# Hemodynamic forces tune the arrest, adhesion and extravasation of circulating tumor cells

**DOI:** 10.1101/183046

**Authors:** Gautier Follain, Naël Osmani, Sofia Azevedo, Guillaume Allio, Luc Mercier, Matthia A. Karreman, Gergely Solecki, Maria Jesus Garcia-Leon, Olivier Lefebvre, Nina Fekonja, Claudia Hille, Vincent Chabannes, Guillaume Dollé, Thibaut Metivet, François Der Hovsepian, Christophe Prudhomme, Angélique Pichot, Nicodème Paul, Raphaël Carapito, Siamak Bahram, Bernhard Ruthensteiner, André Kemmling, Susanne Siemonsen, Tanja Schneider, Jens Fiehler, Markus Glatzel, Frank Winkler, Yannick Schwab, Klaus Pantel, Sébastien Harlepp, Jacky G. Goetz

**Affiliations:** INSERM UMR_S1109, Strasbourg, F-67200, France; Université de Strasbourg, Strasbourg, F-67000, France; LabEx Medalis, Université de Strasbourg, Strasbourg, F-67000, France; Fédération de Médecine Translationnelle de Strasbourg (FMTS), Strasbourg, F-67000, France; Cell Biology and Biophysics Unit, European Molecular Biology Laboratory, Heidelberg, 69117, Germany; Department of Neurooncology, University Hospital Heidelberg, Heidelberg, 69120, Germany and Clinical Cooperation Unit Neurooncology, German Cancer Research Center (DKFZ), Heidelberg, 69120, Germany; Institute of Tumor Biology, University Medical Center Hamburg-Eppendorf, Martinistrasse 52, Hamburg, 20246, Germany; LabEx IRMIA, CEMOSIS, Université de Strasbourg, Strasbourg, F-67000 France; Zoologische Staatssammlung München, Munich, 81247, Germany; Department of Neuroradiology, University Medical Center Schleswig-Holstein, Campus Lübeck, 23535 Lübeck, Germany; Department of Diagnostic and Interventional Neuroradiology, University Medical Center Hamburg-Eppendorf, 20246 Hamburg, Germany; Center for Diagnostics, Institute of Neuropathology, University Medical Center Hamburg-Eppendorf, 20246 Hamburg, Germany; CNRS UMR7504, Institut de Physique et Chimie des Matériaux de Strasbourg (IPCMS), Strasbourg, F-67000, France; LabEx NIE, Université de Strasbourg, Strasbourg, F-67000, France

**Keywords:** Blood flow, extravasation, metastasis, endothelial remodeling, circulating tumor cells, cell adhesion, biomechanics, zebrafish

## Abstract

Metastatic seeding is driven by cell-intrinsic and environmental cues, yet the contribution of biomechanics is poorly known. We aim to elucidate the impact of blood flow on the arrest and the extravasation of circulating tumor cells (CTCs) *in vivo*. Using the zebrafish embryo, we show that arrest of CTCs occurs in vessels with favorable flow profiles where flow forces control the adhesion efficacy of CTCs to the endothelium. We biophysically identified the threshold values of flow and adhesion forces allowing successful arrest of CTCs. In addition, flow forces fine-tune tumor cell extravasation by impairing the remodeling properties of the endothelium. Importantly, we also observe endothelial remodeling at arrest sites of CTCs in mouse brain capillaries. Finally, we observed that human supratentorial brain metastases preferably develop in areas with low perfusion. Altogether, these results demonstrate that hemodynamic profiles at metastatic sites regulate key steps of extravasation preceding metastatic outgrowth.

## INTRODUCTION

Metastatic progression is a complex process resulting in the formation of lethal secondary tumors at distance of its origin^1^. Metastatic cancer cells disseminate very efficiently throughout the body upon intravasation in the blood circulation. Recent work on breast cancer suggests that about 80% of metastases originate from early disseminated cancer cells^2,3^. Once in the blood stream, circulating tumor cells (CTCs) may find a location favoring arrest and stable adhesion before extravasating, and avoiding the hostile shear forces^4,5^. After extravasation, metastatic cells either remain dormant^6^ or grow successfully into life-threatening secondary tumors^7^. Although multiple mechanisms have been postulated for successful extravasation and outgrowth of metastatic cells^7–10^, there are only few insights on the role played by mechanical cues encountered in the blood, the main route for hematogenous metastatic dissemination.

Biomechanical forces are known to have a major impact on metastasis progression. For example, tumor cells sense and respond to stiffening of the surrounding stroma by increasing their invasive potential^11–13^. High extravascular stress caused by tumor growth^14,15^ and interstitial fluid pressure^16^ leads to vascular compression that impairs perfusion and eventually promotes tumor progression, immunosuppression, and treatment resistance. Locally, invading tumor cells need to overcome physical tissue constraints by cellular and nuclear deformability^17,18^, possibly inducing nuclear envelope rupture and DNA damage^19^ leading eventually to inheritable genomic instability^20^. Overall, while the impact of biomechanics on tumor growth and invasion are mechanistically relatively well understood, the *in vivo* mechanisms driving survival, arrest and successful extravasation of CTCs, preceding metastatic growth, remain to be elucidated.

Indeed, very little is known about how CTCs arrest and adhere to the endothelium of small capillaries and leave the blood stream by crossing the vascular wall. While the “seed and soil” concept states that metastasis will occur at sites where the local microenvironment is favorable^21^, the “mechanical” concept argues that arrest and metastasis of CTC occur at sites of optimal flow patterns^22^. CTCs in the blood circulation are subjected to vascular routing^23^, collisions and associations with blood cells^24^, hemodynamic shear forces^25^ and physical constraints imposed by the vessel architecture^7,9^. Only CTCs capable of overcoming or exploiting the deleterious effects of shear forces will eventually arrest, adhere to and exit the vasculature to form a secondary tumor^26^. Nevertheless, a direct contribution of mechanical cues to the arrest and successful extravasation of CTCs has only been poorly studied so far^26^. Therefore, new *in vivo* models, where modeling, visualization and biophysical quantification of the extravasation parameters are easily performed, are of utmost importance for assessing whether biomechanics regulate metastatic extravasation. Here, we aim to address the direct impact of the blood flow on the arrest and extravasation of CTCs *in vivo.* We developed an original experimental approach to measure and modulate blood flow with intravascular injection of CTCs within zebrafish embryos. We observed that blood flow controls the sequential steps of arrest, adhesion and extravasation of CTCs *in vivo.* In parallel, using microfluidics and optical tweezers, we identified the critical adhesion force (80 pN) that CTCs require to initiate adhesion to the endothelium, which rapidly stabilizes under shear flow. This value matches the threshold dragging force measured *in vivo* at extravasation sites. Finally, we used our recently-developed intravital correlative light and electronic microscope (CLEM)^27–29^ to identify endothelial remodeling as one of the major extravasation mechanisms *in vivo*, and that endothelial remodeling scales with flow velocities. Overall our studies demonstrate that blood flow forces at metastatic sites regulate key steps of extravasation preceding metastatic outgrowth.

## RESULTS

### Arrest and adhesion of CTC is favored by permissive flow velocities

In order to test the impact of blood flow on the arrest, adhesion and extravasation of CTCs, we experimentally modeled metastatic seeding in endothelium-labeled zebrafish embryo *(Tg(Fli1a:EGFP))* at 2 days post-fertilization (dpf) by injecting fluorescently-labeled tumor cells in the duct of Cuvier (Figure 1A,B). While metastatic extravasation can be successfully tracked in this model^30^, the zebrafish embryo further allows to combine biophysical characterization and manipulation of blood flow parameters with long-lasting and high-resolution imaging of tumor cells *in vivo.* We first quantified and mapped the position of arrested and stably adherent tumor cells (D2A1) in the zebrafish vasculature and noticed that CTCs preferentially arrested (and extravasated) in the caudal plexus (CP) (Figure 1C). Although arrest and extravasation can be also observed in inter-somitic vessels (ISV) (Figure 1C) and in the brain of zebrafish embryos (Figure 1C, Figure S1A-B, Movie 1), the majority of the tumor cells arrest in the caudal plexus. We exploited the highly stereotyped vasculature of this region for compiling >20 embryos and quantitatively identifying a major hotspot of arrest in this region (Figure 1D), that sits in between the caudal artery and the venous plexus. This was the case for multiple human (Jimt1, 1675, A431), mouse (D2A1, 4T1) and zebrafish (ZMEL) cell lines (Figure S1C). Using fast imaging of the blood flow (100 fps) within the entire zebrafish embryo, combined with PIV (Particle Image Velocimetry) analysis, we observed decreasing flow values in the vascular region that is permissive for the CTC arrest (Figure 1E-F, Movie 2). Accurate dissection of blood flow profiles using PIV analysis showed that flow velocity progressively decreases from the anterior dorsal aorta (Position 1, maximal velocity, v_max_ = 2500 μm/sec, Figure 1E-F), to a minimal flow in its most posterior region (Positions 5 to 6, v_max_ = 500 μm/sec), which we named the arterio-venous junction (AVJ). We have shown in the past that blood flow dissipates along the vascular network of the zebrafish embryo^31^. In addition, the mass conservation implies that ramification of the vessels in the AVJ further contributes to the blood flow decrease. We thus set out to model this phenomenon *in silico* using mathematical simulation of the blood flow in the caudal plexus (see supplementary methods). Simulation experiments reproduced the flow drop observed in the most posterior region of the CP (Figure S1D-H, Movie 3). Besides, to determine whether flow velocity can impact CTCs arrest, we developed an *in vitro* approach to mimic *in vivo* flow profiles in microfluidic channels previously coated with endothelial cells (EC, HUVECs, Figure 1G). We observed that adhesion of CTCs to the endothelial layer was favored by reduced flow profiles (peak velocities of 100 to 400 μm/sec) (Figure 1G), similar to those measured *in vivo* in the AVJ (Figure 1D-E). Using higher flow profiles that mimic flow values obtained in the anterior DA prevented efficient adhesion of CTCs to the endothelial layer (Figure 1G), as observed *in vivo* where no adhesion could be observed in the DA between position 1 and 4 (Figure 1D-E). Adhesion efficacy (and forces) of CTCs was not affected by temperature (28 vs 37°C, Fig.S2B,C). Taken together, these data suggest that reduced flow profiles are permissive for stable adhesion of CTCs to the endothelium, and that the threshold velocity value for efficient adhesion of CTCs ranges from 400 to 600 μm/sec.

**Figure 1:**
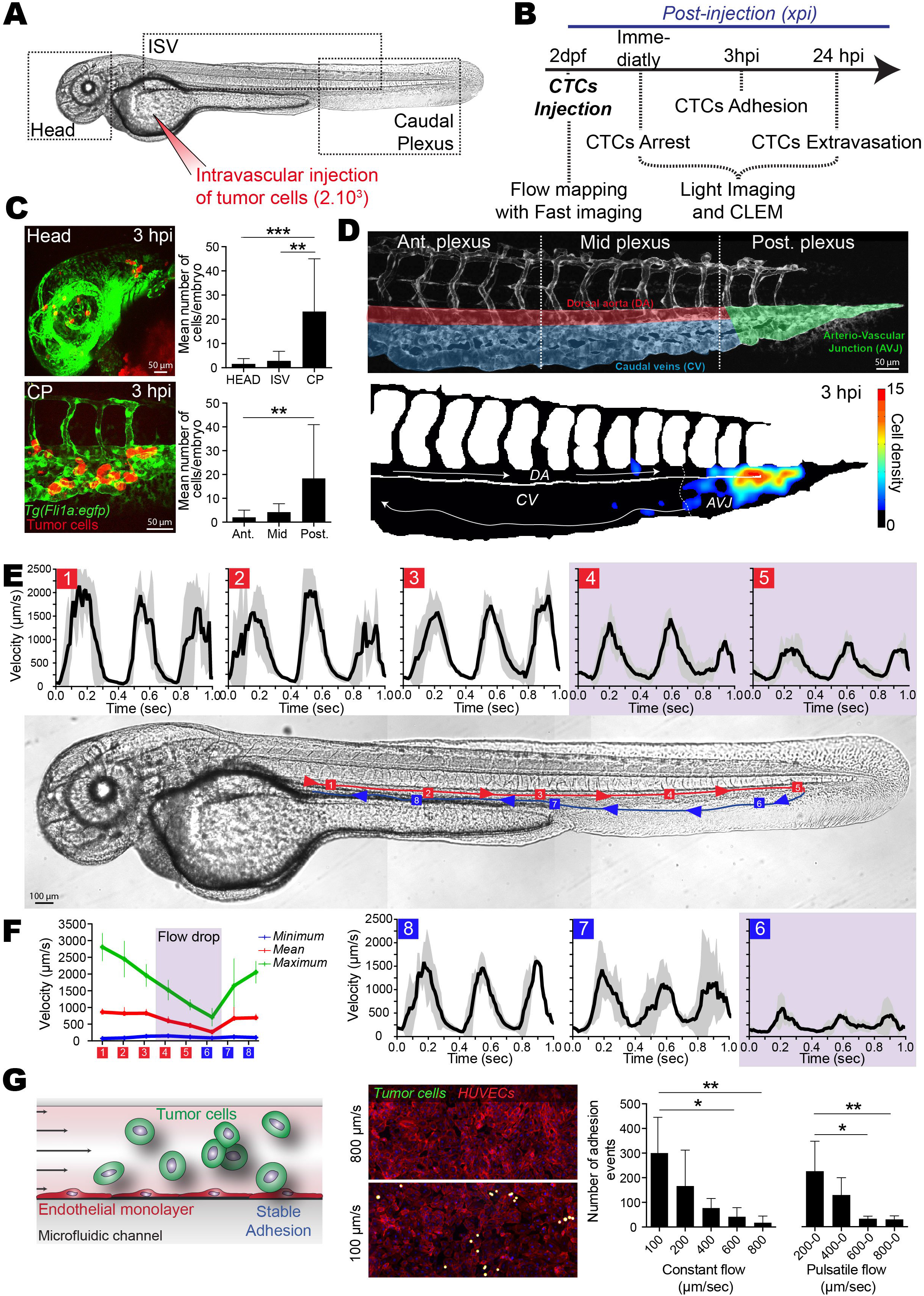
Arrest and stable adhesion of CTCs are favored by permissive blood flow profiles in the zebrafish embryo. (A-B) Experimental design and workflow. xpi: minutes or hours post-injection. (C) Representative images *(Head, head region; CP,* Caudal Plexus) and quantification of arrested CTCs (red) in the vasculature (green) of the zebrafish embryo at 3 hpi (See also Movie 1). Ant., Mid and Post.: Anterior, mid and posterior plexus as depicted in D. (D) High-magnification image of the vascular caudal plexus, the associated regions, and heatmap of arrested CTCs at 3 hpi (n=11 embryos). *AVJ:* arterio-venous junction. *DA:* Dorsal aorta. CV: Caudal veins. Arrows indicate blood flow direction. (E) Blood flow velocity measurements (PIV) in the indicated region (red and blue squares 1 to 8) of the zebrafish embryo. Arrows indicate blood flow direction (See also Movies 2&3). (F) Minimum, maximum and mean values of the blood flow velocity are plotted over the 8 different regions (n=3 embryos). (G) Experimental set-up, representative images and quantification of the microfluidic approach. CTCs (green) are perfused over a monolayer of HUVECs (ECs, red) and adhesion is quantified (n=5 to 6 independent channel per conditions). Pulsatile flow corresponds to 0.3 sec perfusion/0.3 sec stop of the peristatique pump.

### Permissive flow profiles promote stable adhesion of CTCs to the endothelium

Adhesion of CTCs to the endothelium is an important feature that precedes their extravasation^32^. Furthermore, mechanical constraints imposed by cell size and vessel topology likely favor the initial arrest of CTCs^7,33,34^. We set out to test whether such features also contribute to arrest and adhesion in the zebrafish embryo. Our models of CTCs are composed of multiple human (Jimt1, 1675, A431), mouse (D2A1, 4T1) and zebrafish (ZMEL) cell lines whose mean diameters range from 10,23 to 5,25 μm (Fig.2A). We validated these models by comparing their values to average diameters of human CTCs isolated from breast and lung cancer patients (ranging from 7 to 29 μm) (Fig.2A). We then accurately measured mean vessel diameters of the caudal plexus of the zebrafish embryo (Movie 4, Fig.2B). We noticed that diameters of the highly perfused vessels (red, Fig.2B) each displayed a minimal value that exceeds the maximum size for our CTC models (10,23 μm, Fig.2B). When injected in the embryo, our CTCs mostly arrested in the flow-permissive hotspot (AVJ) suggesting that flow drop significantly impacts the arrest of CTCs (Fig.2C). Interestingly, when we correlated the position of arrested CTCs within the vasculature to the vessel size, we observed that 49% of them were located in vessels with diameters <10,23 μm (Fig.2D). These cells could have been trapped in low-size vessels or crawled, upon arrest, in these vascular regions before extravasating. Indeed, we have observed that intravascular CTCs are capable of efficiently crawling on the vessel wall, suggesting that CTCs establish firm adhesions with the endothelium (Movie 5). Moreover, stiff 10-μm polystyrene beads circulate continuously when injected in the embryo, suggesting that size restriction, although it does participate, is not a major determinant of cell arrest (Fig.S2A). These observations led us to test the role of adhesion molecules in driving successful, flow-dependent arrest of CTCs. Thus, we took advantage of our *in vitro* microfluidic approach and used the optical tweezers (OT) technology to trap and stick CTCs to the EC monolayer. Doing so, we identified an average value of 80 pN for the very early adhesion forces (less than 1 min after adhesion of the CTC to the endothelial layer) required for the attachment of CTCs to ECs (Figure S2C, Movie 6). Interestingly, applying the Stokes law to measure the correspondence between flow and force intensity, we noted that a value of 80 pN represents an average flow value of 450 μm/sec, which agrees with threshold flow values that we measured both *in vivo* and *in vitro* (Figure 1). Thus, CTCs establish very quickly adhesion forces that allow them to sustain flow velocities of 450 μm/sec. To test the role of adhesion molecules, we targeted ß1 integrins (ITGB1), known to play a central role in tumor metastasis^35^ and promote stable adhesion of CTC to the endothelium before extravasation^36,37^. We determined whether compromising stable adhesion mediated by ß1 integrins would affect the ability of CTC to arrest under permissive flows. We depleted ITGB1 in our cells using a siRNA approach (Figure 2E) and assessed their stable adhesion 3hpi using a heatmapping procedure over several embryos. Interestingly, while the adhesion hotspot in the AVJ is conserved, the overall number of adhesion event is significantly reduced (Figure 2E). Similar observations were made in our microfluidic system where only 20% of ITGB1-depleted CTCs stably adhere to the EC monolayer using the same perfusion parameters (Figure 2E). Comparable values were obtained when adhesion (or detachment) of perfused CTCs to (from) ECs was forced using OT in microfluidic channels (Figure S2D, Movie 7) indicating, that permissive flow profiles, combined with proper adhesion forces, allow stable adhesion of CTCs to the endothelium.

**Figure 2:**
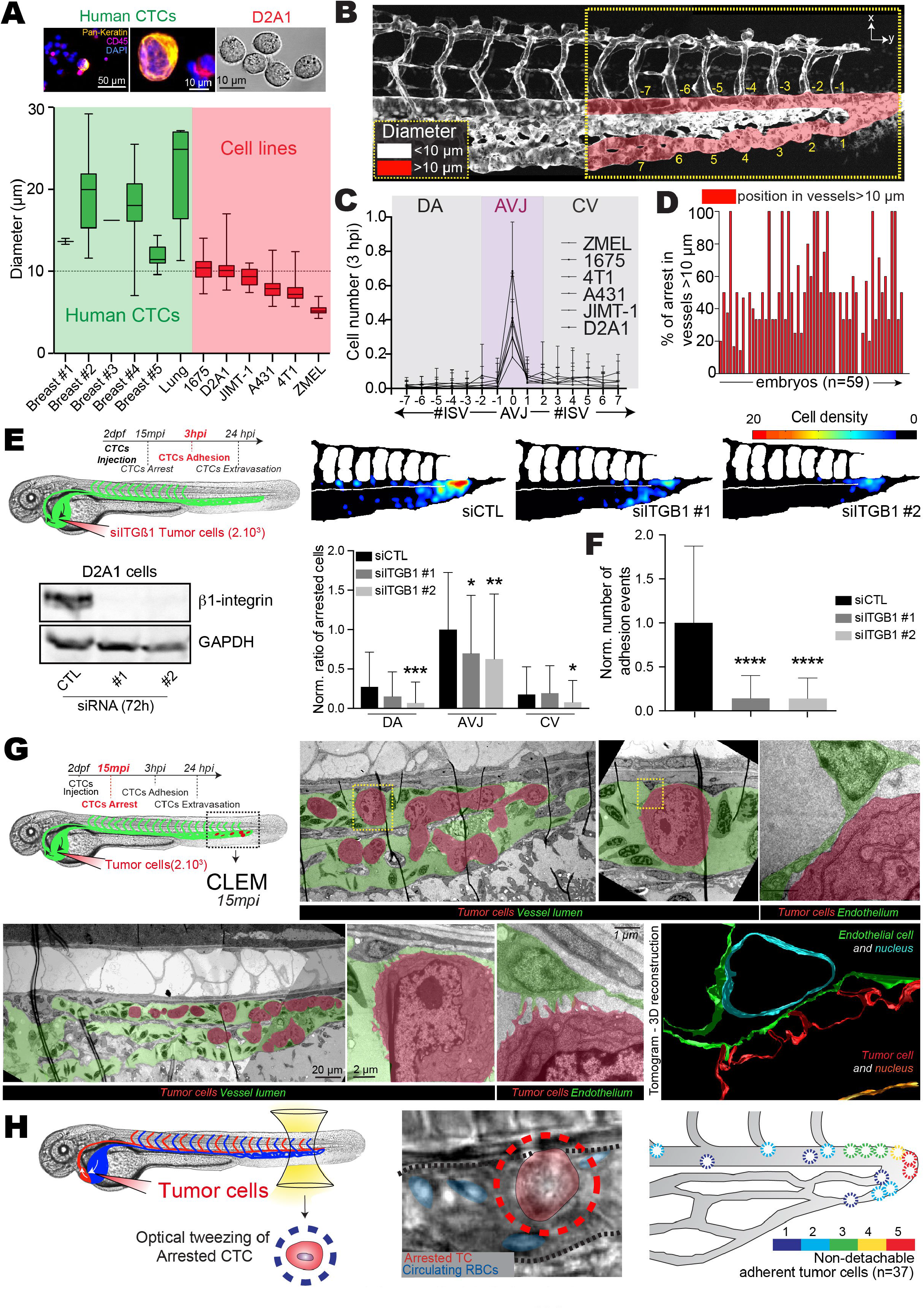
Arrest of CTCs depends on early cellular adhesion forces. (A)Quantification of the diameters of human CTCs (green, five breast cancer patients and one lung cancer patient) and 1675, D2A1, JIMT-1, A431, 4T1 and ZMEL1 cells (red) in suspension. Representative images of human CTCs and D2A1 are also provided. Human CTCs are labeled with Pan-Keratin (CTC marker), CD45 and DAPI (nucleus). (B) Representative confocal Z-projection image of the caudal plexus. Red color shows highly-perfused vessels with diameters over 10 μm (see also Movie 4). (C) Quantification of the number of arrested cells per ISV in the yellow dashed region depicted in B. (D) Quantification of the repartition of arrested cells in the plexus of 59 embryos, between the regions of bigger (red) and smaller (white) than 10 μm vessels in diameter. (E) Experimental workflow and depletion of ITGB1 via siRNA in D2A1 cells: control siRNA (siCTL), and two distinct sequences of the siRNA targeting ITGB1 (siITGB1#1, siITGB1#2). Representative western-blot images are shown. Heatmaps of arrested CTCs after 3hpi are shown for CTCs transfected with siRNAs (n=77, 63, 55, respectively). Quantification of the ratio of arrested CTCs in the indicated region is provided. Norm.: normalized to siCTL AVJ. (F) Graph depicts the number of adhesion events (CTCs to HUVECs) quantified in our microfluidic approach. (See also Movies 6&7). Norm.: normalized to siCTL. (G) Experimental setup of intravital CLEM performed on arrested CTCs only 15 mpi (minutes postinjection). Representative EM images of arrested CTCs (red) in vascular lumen (green) are provided from two different sections (top and bottom). A reconstruction of an Electron Tomogram is provided. (H) Experimental set-up, representative image and quantification of optical tweezing experiments of arrested CTCs in the AVJ (n=37 arrested cells, n>10 embryos) (see also Movie 8).

To further support this hypothesis, we assessed whether adhesion events could be observed *in vivo* using intravital CLEM^38^, which allows to combine live imaging of xenografted CTCs in the zebrafish embryo with Electron Microscopy. Interestingly, CTCs that were arrested in the zebrafish vasculature only 15 mpi displayed fingerlike adhesive contacts with endothelial cells (Figure 2G). In addition, ECs displayed long protrusions when in contact with arrested CTCs (upper panels, Figure 2G). These observations suggest that integrin-dependent adhesion forces between early arrested CTCs and ECs are quickly exceeding stripping forces from the blood flow.

We aimed to further demonstrate the contribution of early adhesion forces to flow-dependent arrest of CTCs and we performed OT *in vivo.* Although OT can very efficiently trap circulating red blood cells (RBCs) in the vasculature of living zebrafish embryo (Movie 8) and detach adhered CTCs from ECs *in vitro* (Figure S3B), OT was inefficient for detaching arrested CTCs *in vivo* (Figure 2H, Movie 8). The inability to detach >35 arrested CTCs with OT, demonstrates that early adhesion forces rapidly exceed 200 pN (which is the technical limit of our OT set-up). Altogether, these results suggest that low flow forces (~80pN) enable the arrest and stable adhesion of CTCs *in vivo.*

### Pharmacological tuning of hemodynamic forces modulates pacemaker activity

Since arrest of CTCs occurs at site of permissive flow patterns, we investigated whether tuning flow forces would impact arrest efficacy. We first tuned the zebrafish pacemaker activity and subsequent flow forces using pharmacological treatments. We selected lidocain^39^ and Isobutylmethylxanthine (IBMX)^40^ to decrease or increase the pacemaker activity (PMA), respectively (Figure 3A,G). Upon treatment, we assessed cardiac PMA and measured an average decrease and increase of 20% for lidocain and IBMX respectively. Using fast imaging combined with PIV analysis we determined the resulting velocities in 3 positions of the DA for several embryos (ISV 1, 4 and 8) and observed that lowering or increasing PMA with lidocain and IBMX, respectively, significantly altered flow profiles (Figure 3B,H, Movie 9). In brief, while lidocain treatments led to lower velocities with longer times under 400 μm/sec (Fig.3B,S3A), IBMX significantly increased the maximum velocities of flow pulses and decreased the overall duration of the flow under 400 μm/sec (Fig.3H, S3A). We confirmed the impact of the two drugs on flow profiles using *in silico* 3D flow simulation (Fig.S3C-F, Movie 10). We further assessed the impact of tuning PMA on hemodynamic forces by trapping RBCs using OT in the AVJ region *in vivo* (Figure S3B, Movie 11). We measured the forces exerted on trapped RBCs, both at the vessel wall and in its center, and extracted the corresponding flow profile based on the Poiseuille law for each condition (Figure S3C). While lidocain significantly reduced flow forces in the center of the vessel, IBMX increased flow forces at the vessel wall and in the center (Figure 3C, I). Importantly, before taking on experiments aiming to study the behavior of CTCs in distinct flow profiles, we demonstrated that our pharmacological treatments had no impact on the vasculature architecture and permeability (Figure S4A-D, Movie 12), on the migratory and adhesive properties of tumor cells (TCs) *in vitro* (Fig.S4E-G) as well as on adhesion efficacies in our microfluidic approach (Fig.S4I).

**Figure 3:**
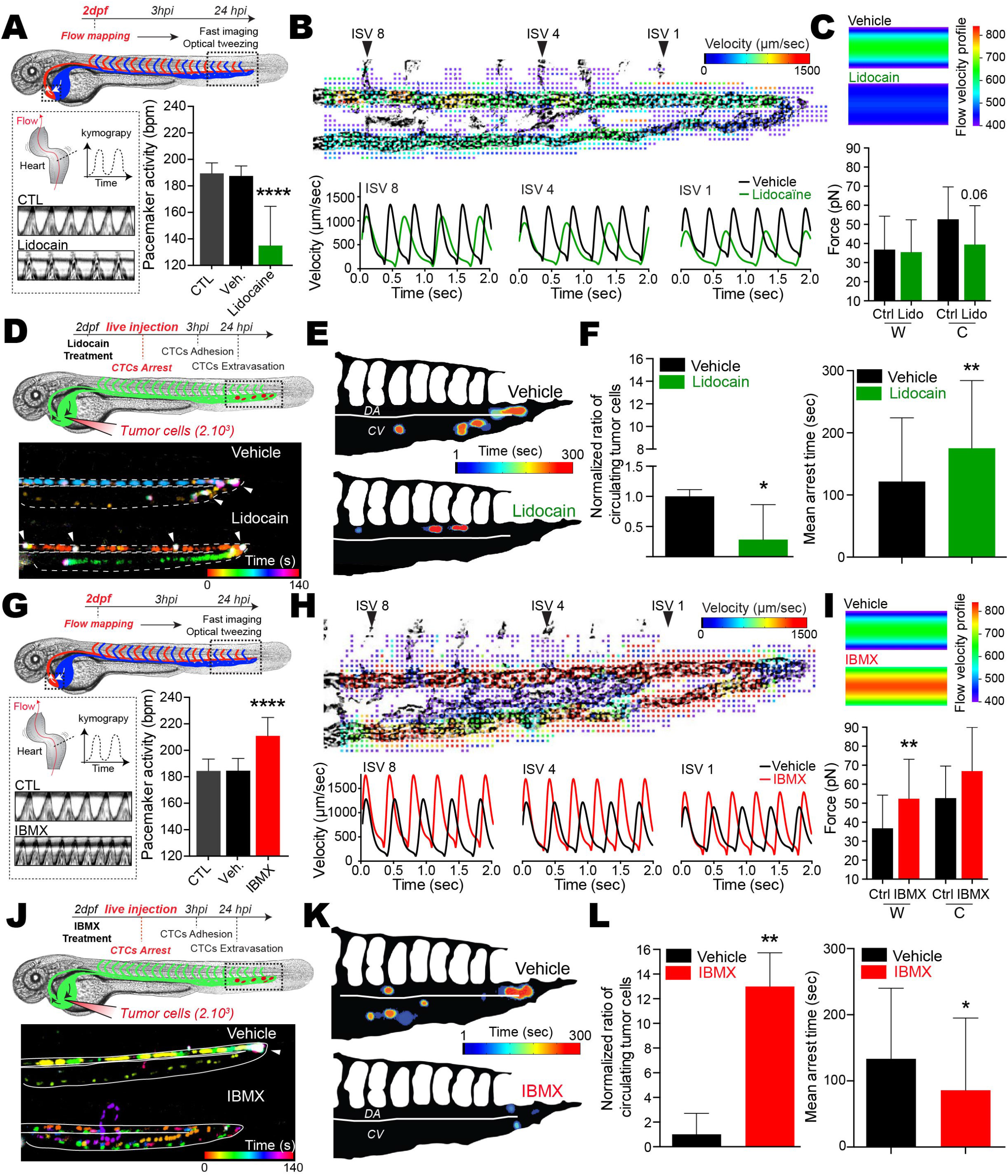
Modulating blood flow forces regulates arrest of CTCs. Zebrafish embryos were treated with vehicles, lidocain (A-F) or IBMX (G-L). (A,G) Experimental workflow. Heart PMA is quantified for each condition. CTL: Breeding water. An additional kymograph analysis is provided. (B,H) (top) Images extracted from fast imaging and PIV analysis of the blood flow in the caudal plexus (region boxed in A,G) (see also Movies 9&10). (Bottom) Quantification of the flow profiles in the center of the dorsal aorta analyzed at position ISV 8, ISV 4 and ISV1 (n=4/5 embryos per condition). (C,I) Orthogonal flow velocity profile and quantification extracted from optical tweezing of RBCs in the dorsal aorta under ISV 1 (W,Wall; C, Center) (See also Movie 11). (D,J) Experimental workflow and representative images of the residency time of CTCs in the caudal plexus over a period of 5 min (See also Movies 13). Arrow heads indicate long lasting arrested cells (white color). (E,K) Spatio-temporal quantification of the residency time of CTCs in the CP using heatmapping on a representative embryo. (F,L) Quantification of the ratio of CTCs as well as the mean arrest time over a period of 5 min for embryos treated with vehicle or the indicated drug. Norm.: normalized to vehicle mean.

### Reduced flow promotes early arrest of CTCs

We then assessed the impact of modulating flow forces on the arrest of CTCs using live and instantaneous intravital imaging upon injection (Figure 3D,J, Movie 13). As expected, CTCs mostly arrest in the AVJ in normal flow profiles (Figure 3D-E, J-K, Vehicle). Interestingly, decreasing flow profiles with lidocain induced arrest of CTCs in anterior regions of the DA, decreased the number of CTCs in the blood flow over a period of 5 minutes post-injection (mpi) and significantly increased their mean arrest time over the imaging period (Figure 3D-F). In contrast, increasing flow profiles with IBMX drastically abrogated the arrest of CTCs in the blood flow over a period of 5 mpi and reduced their mean arrest time (Figure 3J-L). Similar results were obtained when passivated, stiff 10-μm polystyrene beads were injected in control embryos suggesting that inert stiff objects are unlikely to arrest in regions with permissive flow profiles (Fig.S2A). Altogether, these results show that while reduced flow forces enhance the arrest probability of CTCs, increased flow profiles are capable of impeding their early arrest.

### Hemodynamic forces tune the stable intravascular adhesion of CTCs

Because extravasation of CTCs requires stable adhesion to the vessel wall, we investigated whether tuning flow profiles would impact stable adhesion of CTCs to the endothelium *in vivo.* We first assessed the number and location of adhered CTCs 3 hpi in a large number of embryos, making use of our heatmapping protocol and the stereotyped vasculature of the zebrafish caudal plexus (Figure 4A, D). While normal flow conditions favored the definitive arrest in the AVJ, decreasing flow forces with lidocain stimulated the stable adhesion of CTCs in more anterior regions of the DA and reduced the proportion of CTCs arresting in the CV (Figure 4A-C). On the contrary, increasing flow forces with IBMX impaired the stable arrest in the DA and AVJ regions, and shifted definitive arrest towards venous regions of the caudal plexus (Figure 4D-F). Thus, arrest of CTCs mostly occurs in regions with permissive flow profiles, which are shifted towards anterior regions in the DA in lidocaine-treated embryos, and towards posterior regions in the CV in IBMX-treated embryos. We further consolidated this observation by using two additional drugs known to decrease (nifedipin^41^) and increase (norepinephrine^40^) PMA, with similar outcome without perturbing the properties of tumor cells (Figure S4,5A-H). Altogether, these data suggest that flow forces finely tune the adhesive capacity of CTCs, independently of the vessel architecture.

**Figure 4:**
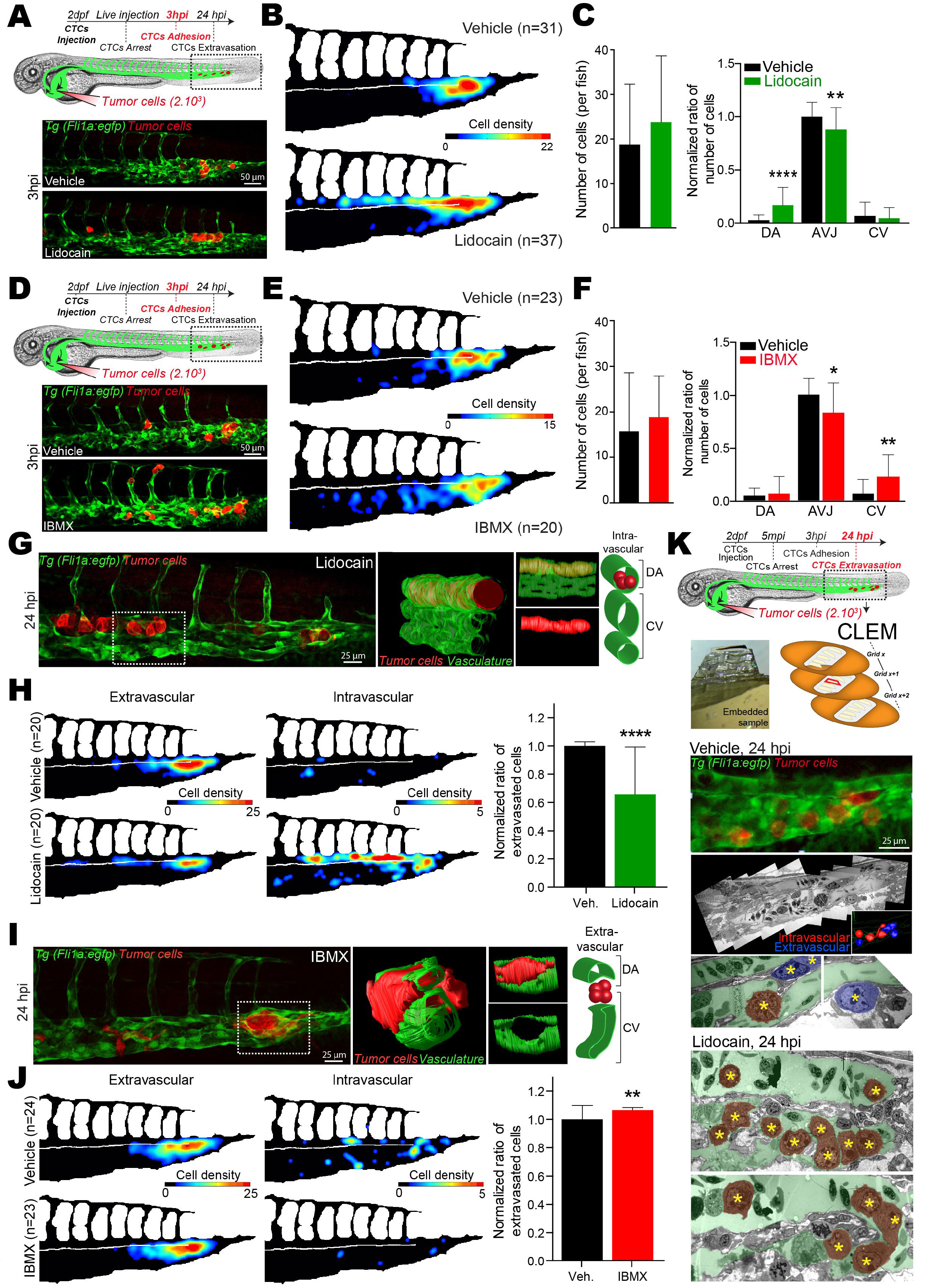
Modulating blood flow forces regulates stable adhesion and extravasation of CTCs. Zebrafish embryos were treated with vehicles, lidocain (A-C and G-H) or IBMX (D-F and I-J). (A,D) Experimental workflow and representative images of arrested CTCs (red) in the caudal plexus (green) of vehicle- or drug-treated embryos. (B,E) Quantification of the number and location of arrested CTCs in the caudal plexus of 3hpi vehicle- or drug-treated embryos, through heatmapping. (C,F) Quantification of the number of arrested CTCs per embryo as well as the ratio of arrested CTCs per region in vehicle- or drug-treated embryos, 3 hpi. Data normalized to vehicle AVJ mean ratio. (G,I) Representative images of TCs (red) in the caudal plexus (green) of 24 hpi vehicle- or drug-treated embryos are shown. 3D reconstruction and scheme of the boxed region is provided. (H,J) Quantification of the number and location of intra- and extravascular TCs in the caudal plexus of 24hpi vehicle- or drug-treated embryos, through heatmapping and histograms. (K) Experimental workflow and CLEM analysis of 24 hpi vehicle- or lidocain-treated embryos is used for further assessing the vascular location of TCs (yellow stars) intravascular (red) or extravascular (blue) (See also Movie 14).

### Increased flow promotes extravasation of CTCs

Once stably attached to the vessel wall, CTCs need to withstand the shear forces and undergo extravasation for metastatic outgrowth. We thus further investigated whether flow forces could impact the extravasation abilities of arrested CTCs and assessed the location and ratio of extravascular cells 24 hpi. Most of the arrested CTCs eventually undergo extravasation in normal flow conditions, while impairing flow forces with lidocain drastically reduced the number of extravascular cells. More than 60% of cells remained fully intravascular, in particular along anterior regions of the DA where CTCs mostly arrested in reduced flow conditions (Figure 4G-H). Impairing flow forces with lidocain only once CTCs were arrested perturbed their extravasation equally (Fig.S4H). Using our CLEM protocol, we confirmed that the TCs were extravascular in normal flow conditions (Figure 4K, Movie 14). Similar experiments conducted on lidocain-treated embryos showed that the vast majority of the cells remain indeed fully intravascular (Figure 4G,H,K). In contrast, increasing flow forces with IBMX further increased the ratio of extravascular cells (close to 100%) (Figure 4I,J), and favored the formation of micrometastasis foci that were surrounded by the local vasculature (Figure 4I). Overall, these results show that blood flow forces enhance the extravasation abilities of CTCs.

### Flow-dependent endothelial remodeling drives extravasation of CTCs

We noticed that ECs could react very quickly to the presence of arrested CTCs (Figure 2G) and that extravasation of CTCs induces massive remodeling of the local vasculature (Figure 4I). In addition, our drugs had no effect on the behavior of the TCs (Figure S4-5), which prompted us to investigate whether flow forces could impact extravasation via ECs. We thus set up long-lasting (15h) intravital 3D confocal recordings of the behavior of TCs during extravasation (Figure 5A, Movie 15). Unexpectedly, image analysis revealed that a majority of CTCs extravasated by indoctrinating the local vasculature rather than by actively transmigrating through the vessel wall by diapedesis, which is nevertheless also observed (Figure 5B-D). Indeed, ECs remodeled around the arrested CTCs by protruding and establishing tight connections with other ECs. During this process, endothelial cells engulf single or cluster of arrested CTCs inducing extravasation of CTCs. In order to gain further structural insights into this phenomenon, we performed intravital CLEM only 9 hpi, for catching an ongoing extravasation phenomenon (Figure 5E). CLEM analysis revealed that clusters of arrested CTCs are indeed fully surrounded by ECs (Figure 5F-G, Movie 16). High-resolution images performed via electron tomography showed that ECs could establish tight contact around the cluster of arrested CTCs (Figure 5H).

**Figure 5:**
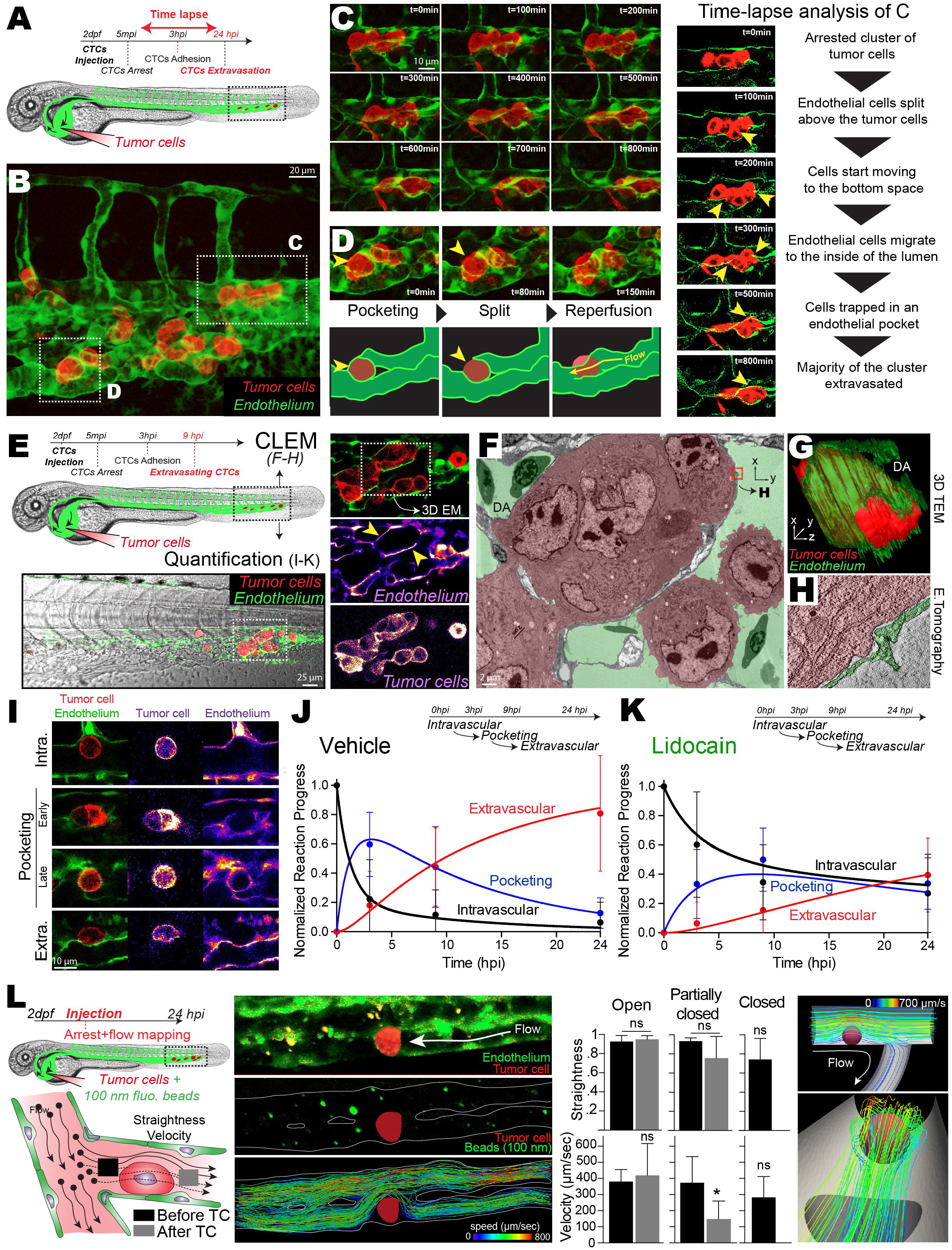
Extravasation of CTCs occurs via flow-dependent endothelial remodeling. (A) Experimental workflow. (B) Extracted Z-projection image from 3D time-lapse analysis of the behavior of arrested TCs (red) in the caudal plexus vasculature (green) over a period of 15h (See also Movie 15). (C) Multiple sequential images, over a period of 800 min, of the region boxed in B are displayed and commented (yellow arrowheads point to the location of the associated comment (left panel)). (D) Multiple sequential images, over a period of 150 min, of the region boxed in B are displayed and commented (yellow arrowheads point to the location of the associated comment). (E) Experimental workflow and CLEM analysis of a 9hpi vehicle-treated embryo. Extracted images of confocal analysis of extravasating TCs (red) at 9hpi is performed. (F) TEM image of the region of interest boxed in E, retrieved in the electron microscopy volume. A representative section is shown and color-coded for TCs (red) and vascular lumen (green) (see also Movie 16). (G) 3D reconstruction of the serial section TEM performed over the entire ROI, depicting TCs (red) and associated ECs (green) (See also Movie 16). (H) Electron tomography image extracted from 3D reconstruction over the boxed region in F (TC: red, ECs: green) (see also Movie 16). (I) Representative images of TCs (red) and the associated vasculature (green) in 9 hpi embryos are shown. (J-K) Quantification and kinetic analysis of the mean number of events over time for vehicle and lidocain treated embryos. The number of events (intravascular, pocketing and extravascular) was quantified at 3hpi, 9 hpi and 24 hpi. (L) Experimental workflow and idealized representation (left). Embryos are injected with CTCs and 100-nm beads and imaged at high speed, before single particle tracking analysis (middle). Corresponding quantification of tracks’ straightness and velocity in 3 scenarios (open, partially closed, closed) and *in silico* representation of laminar flow around arrested tumor cell (right).

We further examined embryos 3, 9 and 24 hpi and documented the behavior of the associated ECs (pocketing) over a high number of arrested CTCs (Figure 5I), over time, and in dependence of the blood flow profiles. Interestingly, in normal flow conditions, injected CTCs that are fully intravascular are rapidly (≈3hp¡) accompanied by EC pocketing that drives successful extravasation of TCs (Figure 5J). On the contrary, reduced flow significantly delays the pocketing behavior of ECs (≈9hpi) impairing efficient extravasation of intravascular TCs (Figure 5K) suggesting that blood flow drives endothelial remodeling.

Although the flow profiles of the zebrafish embryo, and of blood capillaries, is characterized by very low Reynolds number, and are thus mostly laminar, we next wondered whether arrested TCs would significantly impact resulting flow profiles. We injected 100-nm fluorescent beads in the zebrafish vasculature (in addition to CTCs) and performed instantaneous high-speed confocal microscopy (100 fps) for probing flow profiles at high spatio-temporal resolution (Fig.5L). We performed individual track analysis of flowing beads, before and after the position of arrested TCs. Quantification of the tracks’ straightness (1=straight track) and velocities demonstrate that flow profiles are laminar (straightness~1, velocity~400 μm/s) when reaching arrested TCs. In non-occluded vessels, flow profiles remain laminar, with no impact on flow velocity, as expected from our numerical simulations (Fig.5L, Fig.S6A). Indeed, all the streamlines follow the imposed flow direction and do not follow chaotic streamlines. In partially occluded vessels, flow profiles remain laminar, with a slight impact on flow velocity (Fig.S6B). In occluded vessels, flow profiles reaching arrested CTCs are laminar (straightness~1). In addition, we precisely quantified the position of arrested TCs in relation to the first fully perfused vessel: in average, arrested TCs are only separated by <2 μm from the first flow-bearing vessel, suggesting that endothelial cells close to arrested TCs are still exposed to laminar flow profiles (straightness~1, velocity~300 μm/s) (Fig.S6C). Thus, this analysis demonstrates that arrested TCs only deviate established laminar flows present in the vasculature. When arrested CTCs occlude the vessel, laminar flows remain in very close proximity to endothelial cells facing arrested CTCs.

We then wondered whether this flow-dependent phenomenon could be reproduced *in vitro* and set out a microfluidic experiment where *flow* (with or without TCs) is perfused on a monolayer of endothelial cells (HUVECs). Interestingly, stimulating endothelial cells with a laminar flow of 400 μm/s is sufficient to induce a significant protrusive activity of their dorsal surface, which we probed upon expression of a CAG-EGFP construct tagged to the plasma membrane (Fig.6A, Movie 17). Similarly, the dorsal surface of endothelial cells stimulated over 16h with a laminar flow of 400 μm/s displayed characteristic protrusive structures (filopodia and/or microvilli) (Fig.6B). We then perfused CTCs and probed resulting flow profiles around arrested TCs using 100-nm fluorescent beads and high-speed imaging. As we had observed in the zebrafish embryo, arrested TC do not drastically perturb flow profiles, but rather only deviate established laminar flows where streamlines follow the imposed flow direction and do not follow chaotic motion (Fig.S6C). According to this result, CTCs were perfused over a monolayer of ECs for 10 minutes and incubate 16h under perfusion (or not, laminar flow of 400 μm/s) before assessing the behavior of ECs. We observed that flow stimulated the transmigration of TCs. Interestingly, TCs that transmigrated through the HUVECs monolayer were mostly engulfed by ECs, visualized with confocal and scanning electron microcopy, mimicking the pocketing phenomenon we had observed *in vivo* (Figure 6D-E). As a consequence, TCs that successfully transmigrated in absence of flow were rarely surrounded by ECs suggesting that they crossed the monolayer by diapedesis. Altogether, these results demonstrate that the extravasation of CTCs occurs via indoctrination of the associated endothelium, and that flow forces are essential for maintaining the remodeling abilities of the vasculature.

**Figure 6:**
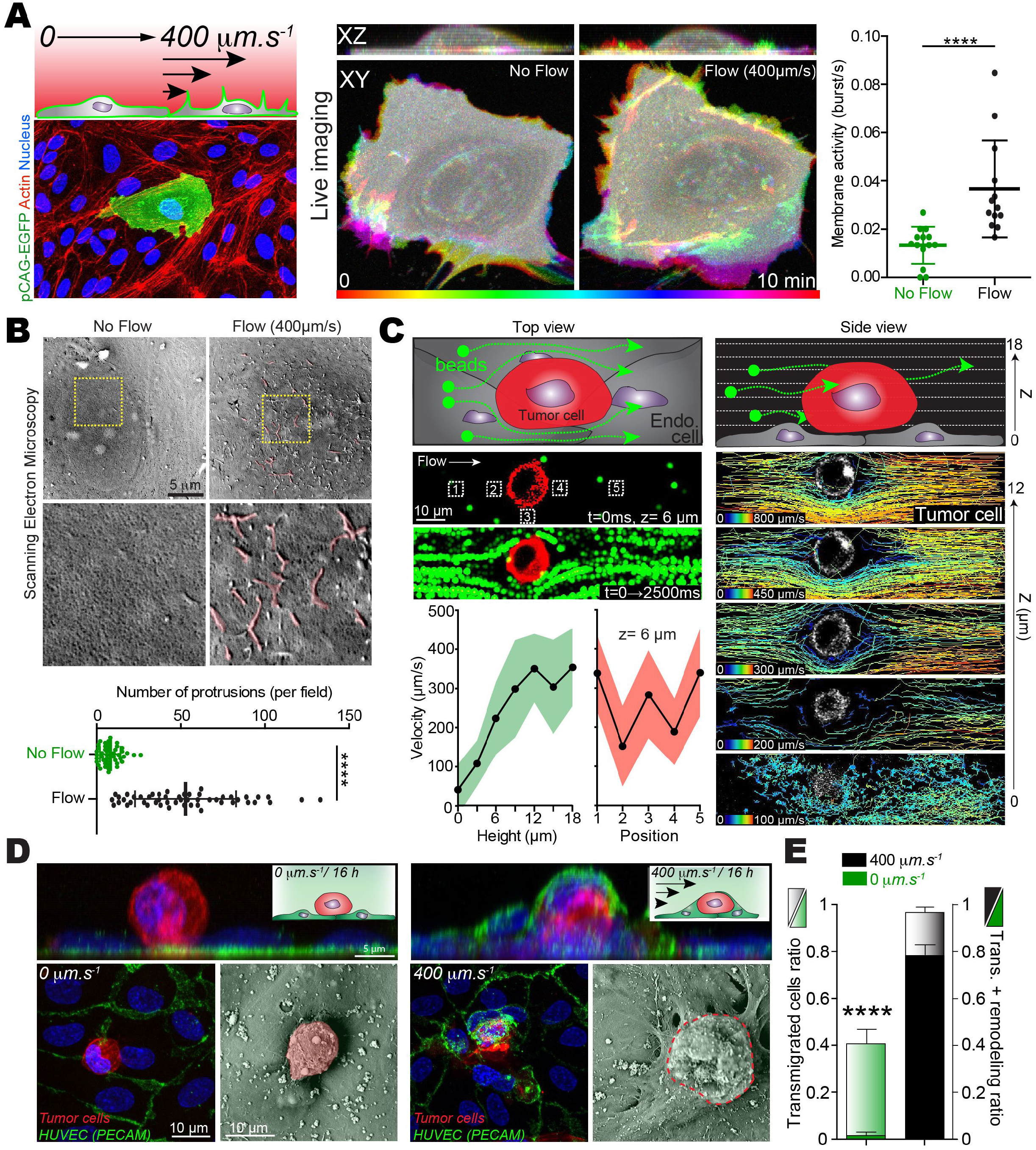
Laminar flow stimulates the protrusive and remodeling activity of endothelial cells. (A) Experimental workflow and representative image of transfected HUVECs. Representative time projection images of pCAG-EGFP expressing HUVECs in flow and no flow conditions, and quantification of the number of protrusion over time. (B) SEM images and quantification showing the number of protrusion in flow vs no flow condition (16h treatment). (C) 100-nm fluorescent beads perfusion over arrested tumor cells in the microfluidic channels. Single particle tracking analysis from high-speed confocal acquisition at different height (Z-step) (right) and quantifications (left). (D) Immunostaining and SEM representative images of TCs (LifeAct, red) arrested on a monolayer of HUVECs (PECAM, green) is shown, in two flow conditions (0 and 400/sec). (E) Quantification of the number of transmigrated TCs and of transmigrated TCs with remodeled HUVECs.

### Micrometastasis develop from mouse brain capillaries with reduced flow profiles

We next investigated whether blood flow forces could explain the location of successful extravasation and micrometastasis formation in another model, the mouse brain. We used our intravital imaging set-up^7^ to quantify the vascular arrest, extravasation, and growth in the mouse brain after intra-cardiac injection of highly metastatic tumor cells (Figure 7A). We analyzed a high number of tumor cells, from several cell lines, that successfully mastered all step of brain metastatic cascade in brain microvessels. We measured the associated blood flow velocities of these vessels, and compared them to flow velocities within blood vessels where arrest and extravasation of CTCs never occurred (Figure 7B). Strikingly, the mean flow velocity in metastasis-proficient vessels was 628±46 μm/sec, which is very close to the permissive flow values we had identified in the zebrafish embryo. In contrast, mean flow velocity of metastasis-deficient vessels was much higher (5880±716 μm/sec). While vessel diameter very likely impacts arrest of CTCs in synergy with permissive flow profiles, additional data suggest that arrested CTC preserve a mild plasma flow, allowing dextran perfusion, while being in close proximity (<10 μm) to vessels perfused with laminar flows (Fig.S7A-D). Together, these data suggest that flow-mediated arrest of CTCs could be a new important and additional determinant of tumor metastasis.

**Figure 7:**
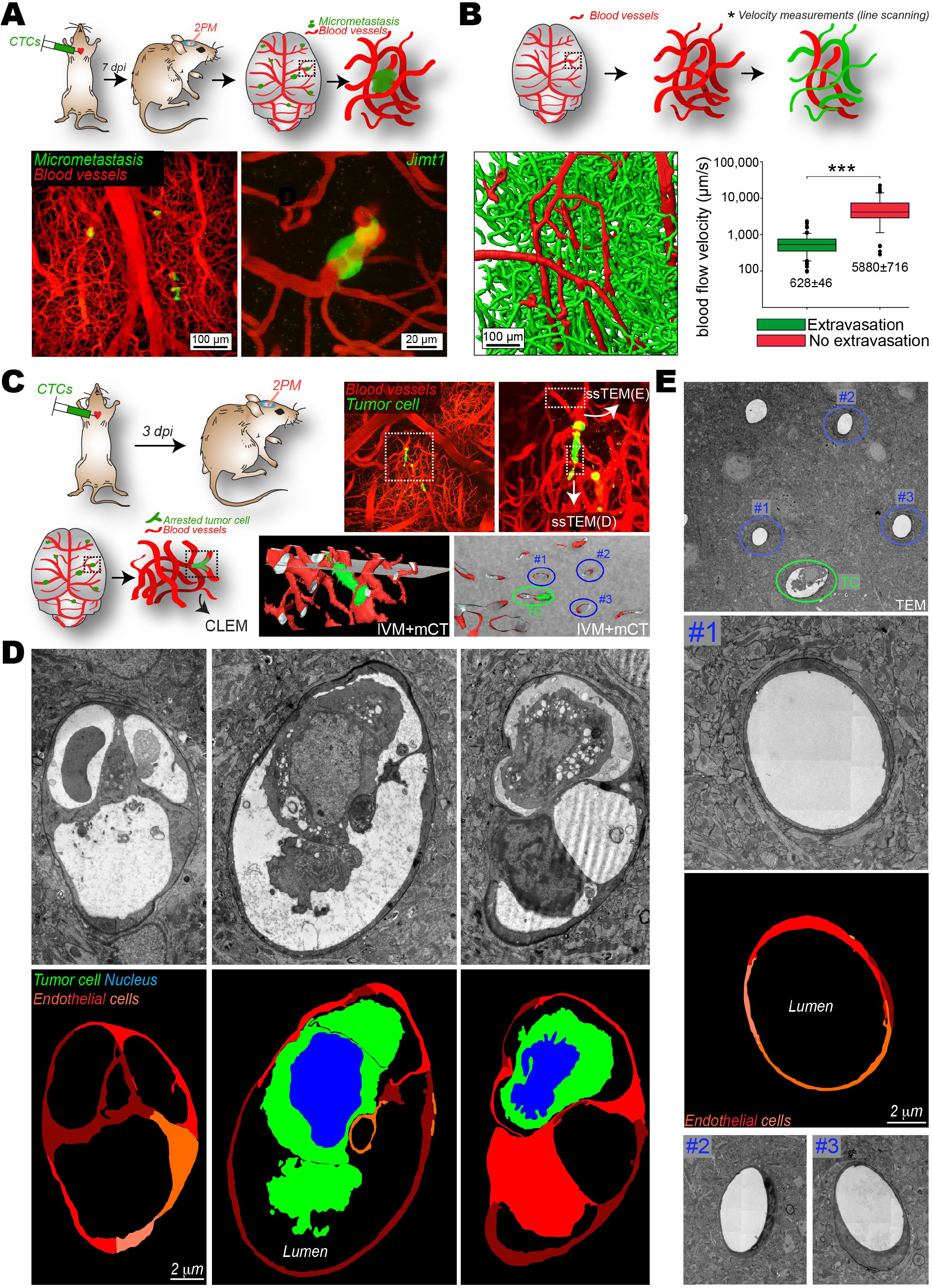
Permissive flow profiles favor formation of micrometastasis in the mouse brain. (A) Experimental workflow and representative images of micrometastases in 7 dpi mice injected with Jimt1-GFP cells (2PM: Two-photon microscopy). (B) Experimental workflow and 3D reconstruction of representative vascular network of the healthy brain. Quantification of the mean blood flow velocity in brain capillary with a diameter of 5 to 10 μm (highlighted in green, where extravasation preferentially takes place) and with higher diameter (highlighted in red, where extravasation is not expected). (C) Experimental workflow of intravital CLEM of 3dpi arrested JIMT1br3 cells. Maximum Z-projections showing an arrested JIMT1br3 cell (GFP, green) and the associated vasculature (TRITC-dextran, red). Merged images of the micro-computed x-ray tomography (mCT) and the intravital imaging (IVM) volume. Vessels containing the TC is circled in green (TC) and dissected using serial section TEM in D, normal neighboring vessels are circled in blue (#1-3) and dissected in E. (D) Serial section TEM images and segmentation of the tumor cell (green), its nucleus (blue) and ECs (red colors) are shown. 3 different z heights are provided. (E) TEM images and segmentation of the ECs (red colors) of normal neighboring vessels are shown.

Similar to the zebrafish embryo experiments, we performed intravital CLEM^28,29^ and investigated whether ECs were also influenced by arrested CTCs in the mouse brain. We thus conducted intracardiac injections of Jimt1 cells and recorded the position of arrested CTCs only 3 dpi, where most of the CTCs are still intravascular (Figure 7C). This enabled us to document, the ultrastructure of the arrested TC (Figure 7C-D), of its associated vessel (Figure 7C-D) and of neighboring vessels where no CTC could be observed (Figure 7C,E). Blood vessels where no arrested CTC could be observed displayed a smooth vascular wall (Figure 7E). However, ECs from the vessel containing an arrested CTC displayed a remodeling phenotype in close proximity to the arrested cell (Figure 7D). Indeed, multiple endothelial cells could be observed protruding within the vessel lumen, establishing contacts with other cells of the vessel wall and enwrapping the arrested tumor cell while maintaining a lumenized vessel. Altogether, these results obtained in the context of brain metastasis (BM) in the mouse suggest that permissive flow profile is a very relevant mechanism driving the arrest of CTCs, leading to micrometastasis growth. In addition, the observation of early vascular remodeling in the presence of arrested CTCs, that are surrounded by plasma and laminar flows (Fig.S7A-C), further suggest an active reaction of the endothelium to arrested cancer cells. Altogether, these observations made in mouse cerebral capillaries speak for very similar mechanisms to what we have observed in zebrafish.

### Blood perfusion controls the location of human brain metastasis

We next investigated whether blood flow perfusion would control the location of BM in humans. For this purpose, we analyzed a randomly selected cohort of 100 patients with BM (n=580) in the supratentorial region of the brain (Fig.8A). The BM of each patient were segmented on contrast-enhanced T1-weighted imaging and the sum mask of all patients (Fig.8B) was co-registered with a computed tomography perfusion atlas of a healthy cohort (Fig.8C). We then performed a voxel-wise comparison of regions containing BM (BM+) and those who did not (BM-) (Fig.8D). Strikingly, BM preferably develop in regions with lower cerebral blood flow (CBF, 51±36-61 vs 58±41-76 ml/100g/min, p<0.001) and higher mean transit time (MTT, 4.6±4.3-5.1 vs 4.5±4.2-5.0 seconds, p<0.001) (Fig.8C,D). These results indicated that occurrence of BM is favored in brain regions with low perfusion, further demonstrating that blood flow patterns are key determinants of metastatic outgrowth.

**Figure 8:**
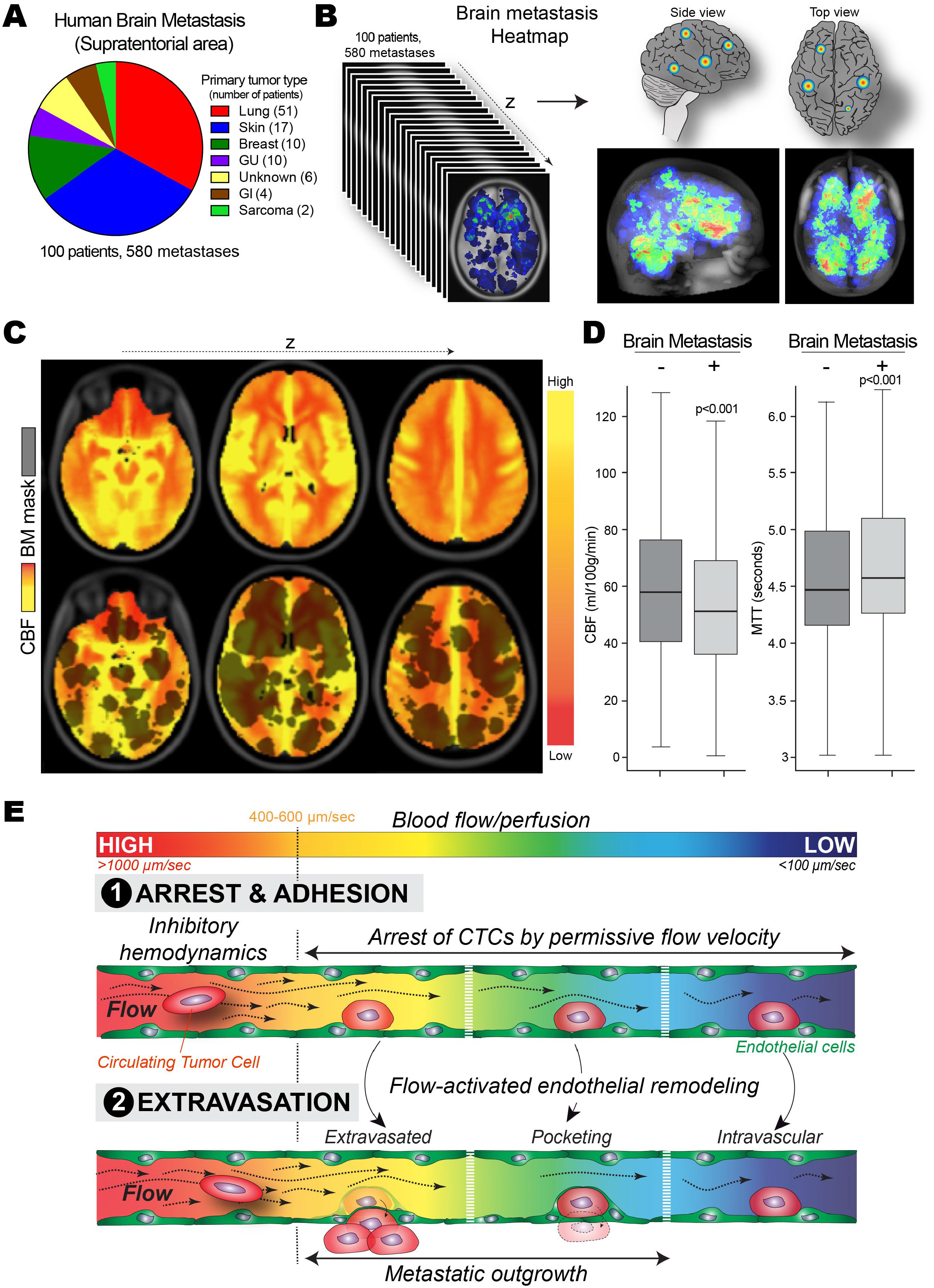
Blood perfusion controls the location of human brain metastases (BM). (A) Number of BM and number of patients are provided for the entire cohort. (B,C) BM of the 100 patients were segmented semi-manually and co-registered on a CT perfusion mask from a control brain template consisting of 107 healthy patients. The cerebral blood flow (CBF) mask is displayed with red indicating a lower CBF and yellow a higher CBF. The cumulative BM is overlaid to the perfusion mask. (D) Mean CBF (cerebral blood flow) and MTT (mean transit time) is displayed for the brain metastases (BM, +) and the corresponding area without any BM (BM, -). (E) Graphical abstract of our study: we identified a threshold of flow velocities that are permissive for the arrest of CTCs, a prerequisite for metastasis formation. Below blood flow velocities of 400-600 μm/s, arrest and adhesion of CTCs to the endothelium is favored. This is the case *in vitro* (microfluidics), in zebrafish embryos and in mouse brain capillaries. Similar results are obtained in humans where a significant reduction of blood perfusion is observed in pro-metastatic regions. In a second step, blood flow is required to stimulate endothelial remodeling and subsequent pro-metastatic extravasation. However, although low flow allows arrest of CTCs, it impairs extravasation of arrested TCs. Thus, pro-metastatic vascular regions are characterized by flow profiles that are sufficiently low (or reduced) to favor the arrest of CTCs, and sufficiently high to stimulate endothelial remodeling.

## DISCUSSION

In this work, we have unraveled for the first time the contribution of flow forces *in vivo* on mediating arrest, adhesion and successful extravasation of CTCs, preceding metastatic outgrowth. Using the multiple advantages of the zebrafish embryo, we have identified a threshold of hemodynamic profiles (400-600 μm/s) that participates to the stable arrest of CTCs in permissive vascular regions, a prerequisite for metastasis formation (Fig.8E). These values were further confirmed *in vitro* and *in silico* using a combination of microfluidic and 3D flow simulation approaches, which were developed upon accurate *in vivo* measurements of hemodynamic forces. We have shown that these permissive flow profiles promote the formation of transient protrusions establishing integrin-dependent adhesion of CTCs to the endothelium. This fast and weak adhesion is then quickly stabilized (less than a minute *in vitro*) to reach adhesion forces exceeding the shear forces generated by such permissive flow values. This process is impaired in regions with high shear values, where forces are significantly higher than the initial adhesion forces CTCs are able to produce. Reduced flow forces favor arrest and metastasis in the zebrafish embryo, but also in mouse brain capillaries. Importantly, we made similar observations in humans where a significant reduction of blood perfusion is observed in pro-metastatic brain regions. Once arrest of CTCs within blood vessels is consolidated, TCs engage in a second step where blood flow remains essential (Fig.8E). Indeed, we have observed that blood flow is required to stimulate endothelial remodeling and subsequent prometastatic extravasation. While reduced flow forces allow the arrest of CTCs, they impede the remodeling potential of ECs, and significantly impair extravasation of arrested TCs. Interestingly, laminar and plasma flows are preserved in regions displaying strong occlusion by arrested tumor cells. Thus, pro-metastatic vascular regions are characterized by intermediate flow profiles that are sufficiently low (or reduced) to favor the arrest of CTCs, and sufficiently high to stimulate endothelial remodeling, which could be a fundamental mechanism driving metastasis formation. While much attention has been given to the contribution of biomechanics to tumor growth and tumor invasion^26,42^, only few evidence suggest that biomechanics could also modulate one of the most important steps of tumor progression, namely extravasation and metastatic outgrowth. However, CTCs are shed in the blood circulation and must survive in a very hostile environment, such as blood shear forces that could induce cell cycle arrest^25^ or even necrotic cell death^5^. In addition, CTCs frequently collide with other components of the blood and the vascular wall, reaching capillary-like vascular regions that appear often occlusive^7,9,36^. Size-mediated arrest of CTCs in capillaries occurs in multiple organs such as the lung, the liver and the brain^43^. Moreover, cellular and nuclear rigidity^18^ and integrity^19,20^ was recently shown to be key drivers of tumor invasion and metastasis. Our work suggests that, in addition to physical occlusion, permissive flows are sufficient to facilitate arrest of CTCs. Selectin-dependent arrest of CTCs has been observed independently of size restriction *in vivo^44^.* In the zebrafish embryo, 50% of arrested TCs did so in vascular regions where diameter of the vessel exceeds diameter of TCs, excluding exclusive occlusion mechanisms. In addition, full occlusion of mouse brain capillaries is rarely observed suggesting that blood flow or physical constraints, although they could impact synergistically, are sufficient to promote the arrest of CTCs. Further work is required to investigate the mutual contribution of cell stiffness/size, vascular architecture and the blood flow as determinant factors in arrest and extravasation of CTCs. Recent work demonstrates that clusters of CTCs, whose arrest would be favored by physical occlusion, are more potent in driving tumor metastasis^45–47^. Besides, we frequently observed a remarkable behavior of arrested CTCs that do not occlude the blood vessel: they crawl counter-flow along the endothelium in perfused vessels and tend to cluster intravascularly before extravasating (Movie 5). This process occurs without compromising blood flow and suggests that TCs establish cell-cell contact that could favor successful extravasation.

A fundamental observation made in our study is that reduced or drops in blood flow favors arrest of CTCs in multiple organisms (zebrafish, mouse and human), preceding metastatic outgrowth. Indeed, we first identified a threshold flow velocity value of 400-600 μm/s that favors the arrest of CTCs in the vasculature of the zebrafish embryo. Using intravital imaging and flow profiling in the mouse brain, we confirmed that brain capillaries perfused with low flow velocities (around 600 μm/s) are favorable to extravasation and outgrowth of TCs. Finally, we analyzed a cohort of 100 patients with brain metastasis and observed that they preferably develop in regions with lower cerebral blood perfusion. Altogether, these results obtained in various organisms indicate that occurrence of metastasis is favored in vascular regions with reduced perfusion, further demonstrating that blood flow patterns are key determinants of metastatic outgrowth in the brain. Interestingly, human brain metastasis had been shown to occur preferentially in vascular border zones as well as in the gray and white matter junction^48^. These regions are located at the most distal part of the arterial tree and display sudden drops in vessel diameter, blood pressure and blood flow, making them very susceptible to infarction and ischemia. Our results further demonstrate that such vascular networks are favorable environments for metastatic extravasation.

Another key observation from our work is that reduced flow promotes arrest of CTCs, by favoring the establishment of intravascular TC-EC adhesions. Indeed, arrest of CTCs can be observed in anterior regions of the DA, when flow values are decreased and depletion of ITGB1 drastically impedes the arrest of CTCs. We identified an initial TC-EC adhesion force of 80 pN, likely mediated by a few integrin molecules. While ITGB1 favors TC protrusion into the subendothelial matrix thereby favoring metastatic outgrowth^49^, integrin-mediated arrest of CTCs could be favored by interaction between ITGB1 and endothelial adhesion molecules, or extracellular matrix molecules localized on the luminal side of the vascular wall. Indeed, *in vitro* data suggest that CTCs hijack the leukocyte adhesion machinery and arrest using α4β1 integrins to bind the VCAM-1 receptor present at the surface of endothelial cells^50^. Furthermore, *in vivo* imaging of liver metastasis has demonstrated that although lumen occlusion by CTCs is rather frequent, it does not account fully for the efficient extravasation as blocking adhesion with ITGB1-blocking antibody significantly impaired stable cell arrest of CTCs^36^. CTCs can escape mechanical entrapment in the lungs^51^ and adhesion-mediated arrest of CTCs has been observed without signs of mechanical entrapment^52^. This suggests that although capillary size can stop CTCs, active adhesion between CTCs and the vascular wall is required for efficient extravasation, and subsequent metastasis. Even if we have not directly visualized this in our experiments, we cannot exclude that CTCs undergo rolling, that could activate ITGB1. Further work will be required to investigate whether arrest of CTCs follows rules that are commonly used by immune cells before extravasation.

An important observation of our work is the route used by arrested CTCs to undergo extravasation. Indeed, in most of the cases, extravasation of CTCs engages massive endothelial remodeling, which encapsulate single or clusters of arrested CTCs. This phenomenon has been observed during extravasation of neutrophils and allows the vessel to reduce vascular permeability^53^. We observed that pocketing or encapsulation of arrested CTCs occurs by the formation of endothelial domes, reminiscent of the ones observed for extravasation of neutrophils^54^ and of leukocytes^55^, that are tightly connected through EC-EC junctions (Figure 6). Recent work performed in zebrafish embryos has shown that stem cells use similar mechanisms to undergo extravasation^56^. Interestingly, using intravital CLEM technology, we observed that vascular regions of arrested TCs in the mouse brain also engage massive endothelial remodeling in vessels with permissive flow profiles, suggesting it could drive extravasation of brain metastatic cells. Altogether, this reinforces the view that CTCs require a very active contribution of the endothelium to perform transmigration of the vascular wall, and challenge the classical idea that CTCs mostly use diapedesis for extravasation^57–59^. Along this line, arrested CTCs have been observed to proliferate attached to the luminal wall of lung blood vessels, leading to intraluminal metastatic foci that eventually passively rupture the endothelial barrier^60^. Recent work suggest that arrested CTCs are also capable of driving TC-mediated necroptosis of EC and subsequent metastasis^10^, further suggesting that the vascular wall is a key contributor to efficient metastatic outgrowth. While endothelium remodeling around arrested CTCs had been observed *in vivo* in the past^30,61^, its importance during metastatic extravasation is still not fully appreciated^62^. Interestingly, past work has shown that cerebral microvasculature is capable of removing blood clots by a process called angiophagy that is strikingly similar to the one we describe here. We now provide evidence that proximal blood flow, which remains laminar, actively regulates such behavior and thereby promoting extravasation of CTCs^63,64^. Indeed, flow forces increase both in vivo pocketing and in vitro endothelial dome formation. As blood flow is known to be key regulator of vascular homeostasis^65,66^ and angiogenesis^38,67^, it is likely that it is required during extravasation by maintaining its remodeling behavior.

In conclusion, we provide the first *in vivo* demonstration that blood flow, which carries tumor-shed CTCs, actively participates in both stable arrest and extravasation of CTCs, preceding metastatic outgrowth. Our work identified pro-metastatic vascular regions that are characterized by laminar flow profiles which are permissive for the the arrest of CTCs, and stimulate endothelial remodeling. Our current work aims to identify the molecular mechanisms that are driven by blood flow to encapsulate arrested CTCs. Altogether, this work suggests that therapies that target endothelial remodeling might be useful to impede extravasation, and subsequent outgrowth, of metastatic cells.

## AUTHOR CONTRIBUTIONS

G.F. performed most of the experiments and analysis, and wrote the paper. N.O. performed the siITGB1 experiments and analysis, and contributed to Fig.3 and Fig.6. S.A. initiated the project, and performed experiments and analysis. G.A. developed the heatmapping protocol and contributed to blood flow analysis, as well as processing of EM images. L.M. and O.L participated to the mouse experiments and statistical analysis. M.K. and Y.S. performed EM of the mouse brain metastasis. G.S. and F.W. performed 2PEM imaging of the mouse brain metastasis. C.H. and K.P. isolated and characterized human CTCs. N.F. performed EM in the zebrafish embryo. M.G.L performed SEM analysis. V.C, G.D., T.M., F.D.H and C.P. developed the flow simulation with input from G.F., S.H. and J.G. A.P, N.P, R.C. and S.B contributed to analysis of the *in vitro* experiments. B.R. and M.K. performed mCT experiments. A.K., S.S., T.S., J.F and K.P. analyzed human brain metastases and blood perfusion patterns on imaging. S.H. supervised the study, performed OT experiments and analysis. J.G. conceived the project, supervised the study, performed experiments and analysis, and wrote the paper with input from G.F., S.H., and N.O.

## ACKNOWLEDGMENTS

We thank all members of the Goetz Lab for helpful discussions We are grateful to Tsukasa SHIBUE (MIT) and Bob WEINBERG (MIT) for providing D2A1 cells, and to Richard WHITE (MSKCC) for providing the ZMEL1 cell line. We are very much grateful to Francesca PERI (EMBL) and Kerstin RICHTER (EMBL) for providing zebrafish embryos. We thank Anita MICHEL (INSERM U949) and Fabienne PROAMER (INSERM U949) from EFS imaging facility for electronic microscopy. We thank Yohan GERBER for help with the Nifedipin and Norepinephrine experiments. We thank Mourad ISMAIL for the fruitful discussions on flow simulation and Marie HOUILLON for participating to the simulation set up. We thank Martin SCHROB (EMBL) for assistance with Electron Tomography. We thank Gertraud OREND, Vincent HYENNE and Michaël POIRIER for critical reading of the manuscript. We thank Raphaël GAUDIN for providing access to the Imaris software. This work has been funded by Plan Cancer (OptoMetaTrap, to J.G. and S.H) and CNRS IMAG’IN (to S.H., J.G. and C.P.) and by institutional funds from INSERM and University of Strasbourg. C.P and V.C were supported by the Center of Modeling and Simulation of Strasbourg (CEMOSIS), ANR MONU-Vivabrain and the Labex IRMIA. G.F. is supported by La Ligue Contre le Cancer. N.O is supported by Plan Cancer. L.M. is supported by an INSERM/Région Alsace Ph.D fellowship. S.A. and G.A. are supported by FRM (Fondation pour la Recherche Médicale). M.A.K. is supported by an EMBL Interdisciplinary Post-doctoral fellowship (EIPOD) under Marie Curie Actions (COFUND).

## STAR Methods

### Contact for reagent and resource sharing

Further information and requests for resources and reagents should be directed to and will be fulfilled by the Lead Contact, Jacky Goetz.

### Experimental model and subject details

#### D2A1 cells (CVCL_0I90). Mouse mammary carcinoma (BALB/c female)

Major information on the D2A1 cell line can be found following this link: http://web.expasy.org/cellosaurus/CVCL_0I90. Culture conditions: 37°/5% CO_2_. DMEM HG with 5% NBCS, 5% FBS, 1% NEAA-MEM, 1% Penstrep. Authentication: Injection in the nipple of mammary gland of BALB/c mice lead to mammary tumor. Cells do not show contamination to mycoplasma.

#### JIMT-1 BR3 cells (CVCL_2077). Human ductal breast carcinoma (Female) highly metastatic in the brain

Major information on the JIMT1 cell line can be found following this link: http://web.expasy.org/cellosaurus/CVCL_2077. Culture condition: 37°/5%CO_2_. DMEM HG with 10% FBS, 1% Penstrep. Authentication: Intracardiac injection in nude mice (NU/NU) lead to cerebral metastasis. Cells do not show contamination to mycoplasma.

#### 1675 cells (WM115, CVCL_0040). Human melanoma (Female)

Major information on the 1675 cell line can be found following this link: https://web.expasy.org/cellosaurus/CVCL_0040. Culture condition: 37°/5%CO_2_. DMEM HG with 10% FBS, 1% Penstrep. Cells do not show contamination to mycoplasma.

#### A431 cells (CRL_1555). Human skin epidermoid carcinoma (Female)

Major information on the A431 cell line can be found following this link: https://www.lgcstandards-atcc.org/products/all/CRL-1555.aspx?geo_country=fr. Culture conditions: 37°/5% CO_2_. DMEM HG with 10% FBS, 1% Penstrep. Cells do not show contamination to mycoplasma.

#### 4T1 cells (CVCL_0125). Mouse mammary gland carcinoma (BALB/c female)

Major information on the 4T1 cell line can be found following this link: https://web.expasy.org/cellosaurus/CVCL_0125. Culture condition: 37°/5%CO_2_. RPMI 1640 with 10% FBS, 1% Penstrep. Authentication: Injection in the nipple of mammary gland of BALB/c mice lead to mammary tumor. Cells do not show contamination to mycoplasma.

#### ZMEL1 cells

Zebrafish melanoma. Culture condition: 28°/5% CO_2_. DMEM HG, 10% FBS, 1% Penstrep. Cells do not show contamination to mycoplasma.

#### Human Umbilical Vein Endothelial Cells (HUVECs)

Primary cells from single donor, commercial vials (PromoCell) amplified before used at P4 for all experiments. Culture condition: 37°/5% CO_2_. ECGM with Supplemental mix (PromoCell) and 1% Penstrep.

#### Zebrafish embryos

*Tg(fli1a:eGFP)* Zebrafish *(Danio rerio)* embryos from a Golden background. Embryos were maintained at 28° in Danieau 0.3X medium (see next section), supplemented with 1-Phenyl-2-thiourea (Sigma-Aldrich) as previously described^38^ after 24 hours post fertilization (hpf). For all Zebrafish experiments, the offspring of one single cross was selected, based on anatomical/developmental good health. Embryos were split randomly between experimental groups. All injection experiments were carried at 48 hpf and imaged between 48 hpf and 72 hpf.

#### Mice

Immunodeficient 8-10 weeks old female Foxn1^nu/nu^ mice (Charles River, Sulzfeld, Germany) were used. Mice were housed at 22°C and have access to water and food at libitum. All work on animals has been carried out in accordance with the German Animal Protection Act after approval by the german ethics committee: Regierungspräsidium, Karlsruhe, Germany. All efforts were made to minimize animal suffering and to reduce the number of animals used. The operation of the chronic cranial window was done as previously described^7,68^. Three weeks after window implantation tumor cells were injected intracardially.

#### Human patients

For the experiments using patient blood collection for CTC size analysis, patient blood samples were acquired in accordance to the World Medical Association Declaration of Helsinki and the guidelines for experimentation with humans by the Chambers of Physicians of the State of Hamburg (“Hamburger Ärztekammer”). All patients gave informed, written consent prior to blood draw.

Human patient data showing the link between brain metastasis (BM) and perfusion pattern were obtained on a single-center cohort. The retrospective study was conducted in compliance with the local ethics committee (Ethik-Kommission der Ärztekammer Hamburg, WF-018/15) with a waiver of informed consent. To collect cases, all MRI studies from 01/2014 to 12/2016 were screened for the presence of untreated malignant intra-axial brain tumors (no previous brain surgery or radiation). In total, 407 patients met the inclusion criteria. From the entire cohort, we randomly selected 100 patients (37 women and 63 men).

## Methods Details

### Cell culture and siRNA-mediated knock-down

D2A1 stably expressing LifeAct-RFP or LifeAct-YPET, kindly provided by Robert A. Weinberg (MIT), were grown as previously described^69^, in DMEM with 4.5 g/l glucose (Dutscher) supplemented with 5% FBS, 5% NBCS, 1% NEAA and 1% penicillin-streptomycin (Gibco). Human Umbilical Vein Endothelial Cells (HUVEC) (PromoCell) were grown in ECGM (PromoCell) supplemented with supplemental mix (PromoCell C-39215) and 1% penicillin-streptomycin (Gibco). To maximize the reproducibility of our experiments, we always used these cells at 4^th^ passage in the microfluidic channels. siRNAs were transfected into D2A1 cells using Lipofectamine RNAiMAX (Invitrogen) following the manufacturer’s instructions. Experiments were performed between 72 h and 96h post-transfection. siRNA sense sequences are the following: siCTL: GCA AAT TAT CCG TAA ATC A, siITGB1 #1: CCA CAG AAG UUU ACA UUA A, siITGB1 #2: GUG UGU AGG AAG AGA GAU A.

### Zebrafish handling

*Tg(fli1a:eGFP)* Zebrafish *(Danio rerio)* embryos from a Tübingen background used in the experiments were kindly provided by the group of F. Peri from EMBL (Heidelberg, Germany). Embryos were maintained in Danieau 0.3X medium (17,4 mM NaCl, 0,2 mM KCl, 0,1 mM MgSO_4_, 0,2 mM Ca(NO_3_)_2_) buffered with HEPES 0,15 mM (pH = 7.6), supplemented with 200 μM of 1-Phenyl-2-thiourea (Sigma-Aldrich) to inhibit the melanogenesis, as previously described^38^.

### Pharmacological treatments

Drugs were added in the breeding water (Danieau 0.3X + PTU) of the embryos before mounting and injection. IBMX (3-isobutyl-1-methylxanthin)^40^, lidocain^39^, nifedipin and norepinephrine^40^ (Sigma) were used at 100 μM in DMSO (incubation time: 20h), 640μM in ethanol (inc. time: 2h), 5 μM in DMSO (inc. time: 2h) and 500 μM in water (inc. time: 1h) respectively. In Fig.S4H, treatment was started 1h postinjection to test the impact of reducing flow forces on extravasation.

### High-speed microscopy for pacemaker activity and blood flow profiling

To measure the heart pacemaker activity and the associated blood flow profiles, we used the USB 3.0 uEye IDS CCD camera (IDEX) mounted on a DMIRE2 inverted microscope (Leica) using transmitted light. Heartbeats were acquired at 80 frames per second (fps) and blood flow in the tail region at 200 fps. For the whole-embryo high-speed recordings of the blood flow, acquisitions were done using an Orca Flash 4.0 cMOS camera (Hamamatsu) mounted on an IX-73 inverted microscope equipped with a UPLFLN 10X/0.3 objective (Olympus). Heartbeats were manually counted to compute the pacemaker activity. The blood flow intensity profile over time at each positions in the vasculature was analyzed using an adapted Particle Image Velocimetry (PIV) protocol from ImageJ software that is available online (https://sites.google.com/site/qingzongtseng/piv).

### Intravascular injection of CTCs in the zebrafish embryo

48 hour post-fertilization (hpf) *Tg(Fli1a:eGFP)* embryos were mounted in 0.8% low melting point agarose pad containing 650 μM of tricain (ethyl-3-aminobenzoate-methanesulfonate) to immobilize the embryos. D2A1 LifeAct-RFP cells were injected with a Nanoject microinjector 2 (Drummond) and microforged glass capillaries (25 to 30 μm inner diameter) filled with mineral oil (Sigma). 23nL of a cell suspension at 100.10^6^ cells per ml were injected in the duct of Cuvier of the embryos under the M205 FA stereomicroscope (Leica), as previously described^30^. The same protocol was used when injecting 1675, A431, JIMT1, 4T1 and ZMEL1 cells. Similarly, injection of 10 μm fluorescent beads (Phosphorex) was performed with a similar concentration of beads, pre-treated with Poly-Ethyleneglycol (sigma) at 0,1 mg/ml in PBS for 5 hours.

### Confocal imaging and analysis

Confocal imaging was alternatively performed with an upright TCS SP5 or SP8 confocal microscope with a HC FLUOTAR L 25X/0,95 W VISIR (Leica), or with a SP2 confocal head mounted on a DMIRE2 stage with HC PL APO 20X/0,7 IMM CORR CS objective (Leica). The caudal plexus region (around 50μm width) was imaged with a z-step of less than 1.5 μm for at least 20 embryos per conditions from at least 3 independent experiments. Cells dispersion was manually counted and localized (using the stereotype patterning of ISVs as reference) in the caudal plexus. The data were compiled to generate heatmaps using a custom-made MATLAB plug in (see next section). Post-processing was performed using Imaris (Bitplane), Amira (FEI) and IMOD (Boulder Laboratory, University of Colorado) for segmentation and 3D reconstruction. For the permeability assay, embryos were pharmacologically treated (as described above) and injected in the duct of Cuvier with 18,4 nl of Dextran-TRITC (500kDa), 3 hours before confocal imaging of the caudal plexus. For assessing the endothelial remodeling *in vivo*, cells were sorted in 3 groups (intravascular, pocketing and extravascular) based on confocal imaging. A kinetic analysis for the individual groups, in vehicle- and lidocain-treated embryos, was performed based on a Michaelis-Menten model, where the arrested tumor cells (TC) interact with endothelial cells (EC) that initiate pocketing (P) before undergoing extravasation (Ex):

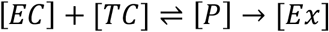

### Heatmaps

The heatmaps were generated using ImageJ (https://imagej.nih.gov/ij/index.html) and MATLAB (MathWorks) softwares. The adhesion/extravasation events are identified for each fish after the analysis of the confocal z-stacks with ImageJ. The position of these events is manually reported on a gray-level support image of a 2.5 days postfertilization zebrafish plexus. Then, all the support images representing each embryo for one condition were put together in an image stack using ImageJ. The stack is read layer by layer in MATLAB and the dots representing the localization are automatically detected with the function Hough circles (Yuan-Liang Tang, Department of Information Management, Chaoyang University of Technology, Taichung, Taiwan) (https://fr.mathworks.com/matlabcentral/fileexchange/22543-detects-multiple-disks--coins--in-an-image-using-hough-transform) using the Circular Hough Transform based algorithm, giving in output the coordinates of the detected dots. Gaussian spots are then created at these coordinates. The amplitude of each Gaussian spot is equal to 1. The different layers of one condition are added each other by making a sum projection, and a black and white mask created with the gray level support image is applied to this sum projection. The gaussian spot amplitudes of each layers are summed to produce the heatmap. The areas of the sum projection where the gaussian spot amplitudes are higher corresponds to high density areas of adhesion/extravasation events. To produce the final heatmap, a custom colormap, inspired by the jet colormap, is applied to the sum projection. The colormap goes from black (no event) to red (high density areas).

### Sample preparation for Electronic microscopy of zebrafish embryos

Correlative Light and Electron Microscopy was performed as previously described^38^. To describe ultra structural characteristics of CTCs and the endothelium in the zebrafish embryo, chosen zebrafish embryos were chemically fixed with 2,5% glutaraldehyde and 4% paraformaldehyde in 0.1M Cacodylate buffer (fish tails cut off in the fixative). Samples were kept in fixative at room temperature for 1-2h and stored in fixative at 4°C overnight or until further processing. Samples were rinsed in 0.1M Cacodylate buffer for 2x5min and post-fixed using 1% OsO4 in 0.1M Cacodylate buffer, for 1h at 4°C. Then, samples were rinsed for 2x10min in 0.1M Cacodylate buffer and secondary post-fixed with 4% water solution of uranyl acetate, 1h at room temperature. Rotation was used at all steps of sample processing. Followed by 5min wash in MiliQ water, the samples were stepwise dehydrated in Ethanol (25%, 50% each 15min, 95%, 3X100% each 20min) and infiltrated in a graded series of Epon (Ethanol/Epon 3/1, 1/1, 1/3, each 45min). Samples were left in absolute Epon (EmBed812) overnight. The following day, samples were placed in a fresh absolute Epon for 1h and polymerized (flat embedded) at 60°C for 24-48h. Once polymerized, most surrounding Epon was cut off using razorblade and samples were mounted on empty Epon blocks (samples flat on the top of the blocks) and left at 60 °C for 24h-48h. Semi-thin sections (200nm) were serially sectioned using ultramicrotome (Leica Ultracut UCT), collected on formvar-coated slot grids and stained with 4% water solution of uranyl acetate for 10min and Reynolds lead citrate for 3min. Semi-thin sections (200nm) were imaged with a CM120 transmission electron microscope (Philips Biotwin) operating at 120 kV. Images were recorded with Veleta 2k x 2k (Olympus-SIS) camera using iTEM software. On the same sections, Electron tomography was performed with a Tecnai F30 Field Emission Gun TEM (FEI) operating at 300 kV and equipped with an Eagle 4K camera (FEI). The F30 was controlled by Tecnai User Interface (PEOUI) and Tecnai Acquisition (TIA) software. Single-axis tomograms were obtained using SerialEM and reconstructed in eTomo, part of the IMOD software package (Boulder Laboratory, University of Colorado).

### Microfluidic experiments

Quantification of the flow-dependent adhesion of CTCs was done using six channels μ-slides VI^0,4^ pre-coated with fibronectin (IBIDI). HUVEC cells were seeded at 21 000 cells per channel (Volume = 30μl). Medium was changed twice a day until they reach maximal confluency (3 to 4 days). D2A1 LifeAct-GFP cells were diluted to maintain the concentration of 150 000 cells/ml of cells between the different speed conditions and perfused using a REGLO Digital MS-2/12 peristaltic pump (Ismatec) and Tygon LMT-55 tubing (IDEX) for 10 minutes before fixation with 4% PFA (Electronic Microscopy Sciences) and stainings (see below). A similar protocol was used for checking the effect of the principal pharmacological treatment (IBMX and Lidocain) and of the temperature (28°C vs 37°C) on the adhesion efficiency of tumor cells.

For endothelial remodeling experiments *in vitro*, two μ-slides I^0.4^ Luer (IBIDI) coated with fibronectin from bovine plasma at 10μg/ml (Sigma F-1141) were used in parallel for each experiment. After seeding the HUVEC cells, one channel was cultured under a flow of 400μm/sec using REGLO pump and Tygon tubbing and for the other channel (no flow condition), medium was changed twice a day. At confluence, D2A1 LifeAct-RFP cells were added at a concentration of 200 000 cells/ml for 10 min. Then, tumor cells were washed using fresh medium and incubated for 16h with or without flow. Z position of the tumor cells relative to the HUVEC monolayer was determined using the piezzo stage of the confocal microscope. A similar protocol was used for checking the effect of the principal pharmacological treatment (IBMX and Lidocain) on the extravasation efficacy of tumor cells.

### Immunofluorescent staining in the microfluidic channels

Cells were fixed using 4% PFA (Electronic Microscopy Sciences), permeabilized with 0.2% Triton-X100 (Sigma) and quenched with 2mg/ml NaBH_4_ (Sigma) 10 min at room temperature before using the following primary antibodies: rat anti-mouse CD29 (9EG7, BD), mouse anti-human CD31 monoclonal primary antibody (MEM-5, Invitrogen). Following secondary antibodies were used: goat anti-rat or mouse coupled with Alexa Fluor 488 (Invitrogen). Cells were mounted using Vectashield (Vector Laboratories). For fluorescent labelling of the HUVEC cells (Fig.1), Alexa Fluor 568 Phalloidin (Life-technologies) and DAPI (Sigma) were used.

### HUVEC transfection and live imaging

HUVECs cell were seeded on fibronectin-coated μSlide I at 100000 cells/channel. The following day, pCAG-EGFP plasmid was nucleofected in HUVEC cells using Nucleofector device and Cell Line Nucleofector Kit V (Lonza) following the manufacturer’s instructions and were seeded into the pre-seeded channels. Live imaging experiments performed 24h later on a DMI6000 equipped with TCS SP5 confocal module and a chamber at 37°C. the HUVEC medium was supplemented with 20 mM Hepes. Briefly, the full volume of single cells was acquired for 10 min sequentially without flow. Then, the flow was turned on at 400 μm/s and the full volume of cells were acquired for 10 min.

### High-speed microscopy for blood flow profiling around arrested tumor cells

48 hour post-fertilization (hpf) *Tg(Fli1a:eGFP)* embryos were injected with CTCs and 100-nm fluorescent beads (Invitrogen), using with a Nanoject microinjector 2 (Drummond) and microforged glass capillaries (25 to 30 μm inner diameter) filled with mineral oil (Sigma). Positively-injected embryos were instantaneously imaged at high-speed (100 fps) using a Leica SP5 confocal microscope equipped with resonant scanner. Such imaging allowed to image arrested tumor cells as well as the displacement of flowing fluorescent beads (100 nm) in their vicinity. We have then developed automated tracking of fluorescent beads (imaged at high-speed) that is followed by single track analysis, before and after the position of arrested TCs. This was performed either using the Imaris (Bitplane) software or Fiji (Trackmate, for in vitro flow proving) Quantification of the tracks’ straightness (net/total distance, 1=straight track) was performed, in addition to flow velocities, for measuring whether flow is laminar or chaotic/turbulent in the presence of arrested tumor cells. A similar approach was conducted *in vitro*.

### Scanning Electron Microscopy on HUVEC and D2A1 in microchannels

HUVECs were seeded in glass coverslips mounted on Ibidi sticky-slide I Luer microchannels, and then cultivated under flow with the addition of D2A1 cells as described in “microfluidic experiments” section. Cells were fixed with glutaraldehyde 2,5% (EMS) in 0.1 M sodium cacodylate buffer (pH7.4), and dehydrated in ethanol-graded series. Once in ethanol 100%, coverslips were detatched from Ibidi sticky chambers. The cell-containing portion of the coverslip was tightly cut with a diamond knife in two pieces, which were irreversibly dehydrated with 1,1,1,3,3,3-hexamethyldizilazane (HDMS, Merck, Millipore)/ethanol-graded series. Next, glass coverslips were mounted onto 25,4mm EM Aluminium Mounts for AMRAY (EMS, Hatfield) by using Leit-C conductive carbon cement (CCC, Plano GmbH, Germany). After proper O/N drying, samples were metalized by platinum vaporization under vacumm up to 12nm (Cressington Sputter Coater 208HR coupled to a Pfeiffer Vacum, Germany). Image acquisition was performed at high-resolution (10kV) and 7000x/ 12000x magnification on a PhenomWorld SEM desktop microscope (Phenom-World B.V, The Netherlands).

### Optical tweezers

Optical tweezing experiments were performed as previously described^31^. Briefly, OT behaves as a picodynanometer where stiffness and the displacement from the steady position can be computed to extract the applied force. Optical tweezing experiments *in vitro* were performed in single channel I^0.4^ Luer μ-slides pre-coated with fibronectin (IBIDI). HUVEC cells were cultured until confluence before the experiment as described above. D2A1 were perfused at 10^4^ cells/ml at a low speed to maximize the cell trapping and avoid collision effects by other CTC. A 1064 nm laser (Navigator 1064-7Y Spectra Physics) was used to generate optical tweezers mounted on a thermostated inverted microscope equipped with a UPLFLN-P 100X/1.3 objective (Olympus). D2A1 cells attached to the HUVEC monolayer were trapped in the beam and moved away from the HUVEC monolayer by displacing a computer-controlled piezzo stage (PI P545 PNano) along the X/Y axis. The fluctuations of the cell in the trap were recorded on a quadrant diode (Thorlabs PDQ30C) and the associated spectral power density allowed the calibration of the trap stiffness. The detachment of D2A1 cells from the HUVEC monolayer was recorded using a Cmos camera (Thorlabs DCC3240C). The movies were analyzed to access the center of mass of the cell in each image. This position compared to the steady position of the cell in the trap is proportional to the force exerted on the cellular adhesion. The rupture force between D2A1 and HUVEC gave rise to a drop in the position of the trapped cell steady state and allowed to access the rupture force.

Optical tweezing experiments in the zebrafish embryo were performed as previously described^31^, with slight modifications: We implemented the setup with a thermoregulated chamber ensuring the embryos to remain at 28°C. We acquired the displacement with the camera at high frame rates 200fps after having established the power spectrum from the photodiode at 20kHz resolution to calibrate the setup.

### Mice experiments and surgical procedures

8-10 weeks old female Nu/Nu mice (Charles River, Sulzfeld, Germany) were used to study the extravasation cascade. All efforts were made to minimize animal suffering and to reduce the number of animals used. The operation of the chronic cranial window was done as previously described^7,68^. Three weeks after window implantation a 100μl cell suspension, containing 500,000 tumor cells (JIMT1br3, A2058br, PC14PE6br, H1), was injected intracardially. *In vivo* imaging was done on day 3 and day 7 after the heart injection. After imaging, the mouse was perfusion-fixed through intracardiac injection with 2.5% glutaraldehyde (Electron Microscopy Sciences, Hatfield, PA) and 2% formaldehyde (Electron Microscopy Sciences) in 0.1 M PHEM buffer (comprising 60 mM PIPES, 25 mM HEPES, 10 mM EGTA and 2 mM MgCl, pH adjusted to 6.9). Following fixation, the regions of interest (ROIs) were imaged again based on the stored stage x,y-coordinates. NIRB was performed with the same laser that had been used for *in vivo* microscopy, tuned to 800 nm wavelength, as previously described^29^. Above the ROIs, at the level of the brain surface, a 150×150 μm^2^ area was scanned in a single focal plane until the NIRB square became clearly visible through emission of autofluorescence in the green channel. Around the ROI, three bigger 300×300 μm^2^ NIRB squares were drawn in non-symmetric positions to facilitate orientation and retrieval of the ROI upon dissection. The brain was removed from the skull and post-fixed by immersion in the same fixative at 4°C overnight. The following day, the fixative was replaced with 0.1 M PHEM buffer, and the brain was stored at 4°C until further processing^29^. All animal procedures were performed in accordance with the institutional laboratory animal research guidelines after approval of the Regierungspräsidium Karlsruhe, Germany (governmental authority).

### *In vivo* multiphoton laser scanning microscopy (MPLSM)

*In vivo* imaging was performed with a Zeiss 7MP microscope (Carl Zeiss Microscopy, Jena, Germany) provided with a Coherent Chameleon UltraII laser (Coherent, Glasgow, UK) with a 500-550 nm and a 575-610 nm band pass filter. With the wavelength of 850 nm GFP and TRITC-dextran were detected. To prevent phototoxic effects, laser power was always kept as low as possible. During the imaging process animals were anaesthetized with a low gas narcosis including 1.5% isoflurane (Baxter, Unterschleißheim, Germany) and 98.5% oxygen (Guttroff, Heidelberg, Germany). During imaging body temperature was kept constantly at 37°C by a heating pad. To acquire angiographies of brain blood vessel, 0.1 ml TRITC-dextran (500 kDa, 10 mg.ml^−1^, Sigma-Aldrich, Munich, Germany) was injected.

### Blood flow velocity measurements and visualization of MPLSM data

The blood flow velocity was measured by a line scan with a minimum length of 10 μm, detecting 2000 events in microvessels. At least 16 randomly chosen vessels were measured per animal. The resulting scan identifies single erythrocytes as angular black lines, where the x-axis is according to the length of the detected distance and the y-axis is the elapsed time of the measurement. The angle is converging more and more to 90° when the cells are static. By knowing the resolution of both parameters and by the measurement of the slope of 30 randomly chosen red blood cells a calculation of the mean velocity is possible, by inversion of the result.

### Western blotting

For western blotting analysis, extracts corresponding to similar cell numbers were loaded on 4-20% polyacrylamide gels (Biorad) and run under denaturing conditions. The following primary antibodies were used: ITGB1 (Millipore AB1952; rabbit), GAPDH (V-18; Goat). HRP-conjugated secondary antibodies were used with ECL (GE Healthcare) for reading using using a PXi system (Syngene). Intensities were normalized over cellular GAPDH levels.

### *In vitro* migration and adhesion assays

For wound healing assay, cells were cultured until confluence in 6-wells plate. The monolayer was scratched with a 20 μl tips and medium was changed with medium supplemented with the corresponding pharmacological treatments. Pictures were taken at time = 0h, 3h, 6h, 9h and 24h. Wound closure was analyze using ImageJ.

For adhesion assay, cells were plated at 20 000 cells/ml in 96-well plate filled with culture medium supplemented with the corresponding pharmacological treatments. After 30 min incubation, wells were washed with PBS, fixed in 4% PFA and stain with 1% violet crystal (VC, Sigma) for 1h at room temperature. VC was washed with PBS and pure DMSO was added to solubilized VC for 30 min under gentle moving. Optical density at 590nm was measured using TriStar^2^ plate reader (Berthold).

### Mathematical modeling and numerical methods for *in silico* experiments

Detailed protocol for the in silico experiments are provided as a stand-alone supplementary file. We conducted numerical simulation for the different pharmacological treatments (IBMX, EtOH, Lidocain, DMSO) as well as for several positions of arrested tumor cells. Each condition requires high performance computing to solve the equation set. We used 48 cores to solve each configuration to simulate 4s of blood flow. The later ensures that the flow is correctly established.

### Patient blood collection for CTC size analysis

Patient blood samples were acquired in accordance to the World Medical Association Declaration of Helsinki and the guidelines for experimentation with humans by the Chambers of Physicians of the State of Hamburg (“Hamburger Ärztekammer”). All patients gave informed, written consent prior to blood draw. Samples were collected from 5 metastatic breast cancer patients into standard 7.5 ml ethylenediaminetetraacetic acid (EDTA) vacutainers and transferred to TransFix^®^ (Cytomark) tubes within 2 h of sample draw. Blood was also collected from a metastatic non-small lung cancer patient directly into either CellSave^®^ (Menarini-Silicon Biosystems), Cell-free DNA BCT^®^ (Streck) or TransFix^®^ preservatives for further analysis. Circulating tumor cells were isolated from whole blood sample by two size based enrichment systems: Both methods have been validated within a large European consortium of 36 partners from academia and industry, CANCER-ID (www.cancer-id.eu). The Parsortix^™^ system (ANGLE plc, UK) enriches tumor cells from a wide variety of blood tubes and has been described in detail in previous publications^70,71^. In short, the device passes blood into a disposable cassette. This cassette narrows down to a critical gap of 6.5 μm at which all larger cells are retained while smaller cells pass through into the waste, thus drastically reducing white blood cell and erythrocyte background. Due to their larger size and higher rigidity, tumor cells are retained at the gap and can be extracted by reversing the flow within the chamber. The enriched cell fraction is harvested into a cytospin funnel and spun down (190 x g, 5 min). Upon drying overnight, samples can either be frozen at − 80 °C or directly stained by immunocytochemistry (ICC). The second size based enrichment system used to process cancer patient samples, was a novel microfluidic filtration device provided by Siemens Healthcare Diagnostics (Elkhart, IN, USA). The system uses a Hamilton STARlet™ robot (Hamilton Company, Reno Nevada) with a specifically developed software allowing for automated enrichment and staining of CTCs from whole blood^72^. Briefly, the blood is passed through a membrane containing pores of 8 μm, enriching for all larger cells. For this, EDTA blood samples were transferred into TransFix^®^ preservation tubes and incubated overnight. The blood was then transferred to the device together with all reagents necessary for ICC staining of CTCs. Cytospins generated following Parsortix^™^ enrichment were stained using fluorescently labeled antibodies. ICC included fixation with 4 *%* PFA, permeabilization with 0.1 *%* Triton X-100 and blocking using serum albumin. Two pan-Keratin antibodies (clone AE1/AE3, eBioScience and clone C11, Cell Signaling) were combined with a CD45 targeting antibody (REA747, Miltenyi Biotec) and DAPI for cell detection. Cytospins were cover slipped using ProLong^®^ Gold antifade reagent (Invitrogen). For the filtration device, we performed a single ICC staining step using a cocktail of fluorescently labeled antibodies, followed by cover slipping with ProLong^®^ Gold antifade reagent (Invitrogen). Tumor cells were detected by targeting Dy550 labelled keratins (Pan CK – clone AE1/AE3, CK8/18 – clone UCD / PR10.11, CK19 – clone A53-B/A2), multiple Dy650 labelled white blood cell markers (CD45 – clone 9.4, CD66b – clone G10F5) and DAPI, all provided directly by Siemens Healthcare Diagnostics. Cytospins and filtration membranes were evaluated and enumerated manually by fluorescence microscopy. CTCs were classified as pankeratin positive, CD45 negative and DAPI positive cells. They were photographed using the AxioVision LE64 microscope software (Zeiss) which allows for measurement of lengths with its respective processing tools.

### Brain metastasis and blood flow perfusion study

MRI was performed using a 1.5 Tesla (Magnetom^®^ Sonata, Siemens Healthcare, Erlangen, Germany; Magnetom^®^ Symphony, Siemens Healthcare, Erlangen, Germany, and Magnetom^®^ Avanto, Siemens Healthcare, Erlangen, Germany) in 94 patients or a 3 Tesla scanner (Magnetom^®^ Skyra, Siemens Healthcare, Erlangen, Germany; Ingenia, Philips Medical Systems, Best, The Netherlands) in 6 patients. Imaging protocol always included axial T1w spin echo with flow compensation and/or three-dimensional T1w gradient echo sequences following weight-adjusted Gadolinium injection (T1w+). If both sequences were acquired the latter one was used for further analysis. Sequence parameters (repetition time, echo time, inversion time, field of view, matrix, pixel size, slice thickness, interslice gap, and number of slices) varied among the different scanners. All BM were subsequently segmented semi-manually using the Analyze Software System 11.0 (Biomedical Imaging Resource, Mayo Clinic, Rochester, MN, USA)^73^. Axial T1w+ MR images of all patients were then automatically co-registered to the Montréal Neurological Institute (MNI) brain by using the FMRIB Software Library (Analysis Group, Oxford, UK) linear registration tool. Correct registration of all T1w+ images and the segmented BM to the MNI brain was secured by visual inspection. Cluster maps comprising all BM of the respective cohort were calculated (Figure 8). Computed tomography was used to build the brain perfusion atlas. Since cancer patients often show an altered cerebral blood flow due to paraneoplastic changes or chemotherapy administration, we decided to use computed tomography (CT) perfusion data from a healthy cohort as reference^74,75^. In brief, 107 patients were triaged by CT perfusion for symptoms of transient ischemic attack but without evidence of ischemia or any perfusion abnormality, infarction or symptoms on follow up, or vascular abnormality as reported elsewhere^76^. Quantitative perfusion maps were obtained for CBF and MTT^77^. All perfusion raw data were processed in a central core-lab on a workstation dedicated for perfusion analysis (Syngo mmwp VE52A with VPCT-Neuro; Siemens Healthcare, Forchheim, Germany) with motion correction and low band temporal noise removal. Non-parenchymal voxels corresponding to bone, vasculature, calcification and cerebrospinal fluid were automatically excluded by adaptive intensity thresholding. Perfusion parameter maps were calculated based on a deconvolution model by least mean squares fitting. All perfusion maps were then affine registered to 1 mm MNI standard space by a precise registration model between the baseline time average of each CT perfusion dataset and a custom CT template in standard space using the FMRIB Software Library 5.0^78^. Mean voxel-wise perfusion parameter maps normalized to standard space were then registered to each individual patient in our study cohort to obtain voxel specific normal perfusion values. Absolute mean cerebral blood flow (CBF in ml/100g brain tissue/min) and mean transit time (MTT in sec) values of all BM were calculated for each voxel of the perfusion mask and each voxel with a BM. Statistical analysis was conducted using IBM SPSS Statistics^®^ software (IBM^®^ 2011, version 20, Armonk, New York, USA) and R (The R Foundation, version 3.3.1. Vienna, Austria). The differences in CBF and MTT between all BM and BM- voxels were compared by the independent t-test. Voxels with more than one BM were weighted according to the number of BM occurring within the voxel. The paired samples t-test was used to compare the perfusion values and the volume of the BM. If not otherwise indicated, data are given as median ± interquartile range.

## Quantification and statistical analysis

### Statistical tests

Statistical analysis of the results obtained during Zebrafish, Mice and microfluidic experiments were performed using the GraphPad Prism program version 5.04. The Shapiro-Wilk normality test was used to confirm the normality of the data. The statistical difference of Gaussian data sets was analyzed using the Student unpaired two-tailed t test, with Welch′s correction in case of unequal variances. For data not following a Gaussian distribution, Kruskal-Wallis and Dunn’s multiple comparisons post-test or the Mann-Whitney test was used. For qualitative data, the Fisher test was used. Illustrations of these statistical analyses are displayed as the mean +/− standard deviation (SD). p-values smaller than 0.05 were considered as significant. *, p<0.05, **, p < 0.01, ***, p < 0.001, ****, p < 0.0001.

Please find bellow, the details related to sample size and data exclusion, replication and randomization for each experiment.

### Zebrafish experiments

The Pacemaker activities of the embryos (**Fig 3A and G, Fig S5A and E**) had been registered during 3 to 5 experiments all along the experimentation as a control of treatment efficiency. Represented sample sizes are: n=60 Veh. Lidocain, n=78 lidocain, n=58 Veh. IBMX, n=65 IBMX, n=11 Veh. nifedipin, n=10 nifedipin, n=10 Veh Norepinephrin, n=10 Norepinephrin.

Blood flow measurement regarding pharmacological treatment using PIV/automated treatment (**Fig S3A, Fig 3B and 3H**) was done on 5 embryos from the same batch, Except for one Veh. EtOH treated embryos that was excluded during analysis, based on imaging focus default.

Blood flow measurement using PIV/automated treatment on ZF whole embryo (**Fig 1E-F**). Sample size = 3 fish. Due to the size of the whole embryo, the imaging time and cost of acquisition are very high, so we decided to limit the sample size.

Optical tweezing of adhered tumor cells (**Fig 2H**) were performed on 37 cells during 4 independent experiments.

Blood flow measurement using optical tweezing (**Fig 3C and I, Fig S3B**) was done over more than 5 embryos per condition from several treatment batch. due to the low throughput of the approach and knowing that the pacemaker activities from both vehicle condition are not significantly different, we decided to pull them as a control situation. Sample sizes for Ctrl / IBMX / Lidocain:

- number of cells in total: 75 / 54 / 54
- trapped at the wall of the DA (Fig 3C): 26 / 22 /19
- trapped in the center of the DA (Fig 3C)”: 23 /7 / 12

All Zebrafish injection experiments at 3, 9 and 24 hours post-injection (**Fig 1C-D, Fig S1C, Fig 2C-E, Fig 4A-J, Fig S4H, Fig 5I-K, Fig S5 B,C S5G,F and S5I**) were repeated at least three times independently on 16 embryos per condition each time. Taking into account, the success rate of injection and imaging time limitation, this lead to the following final numbers:

- 3hpi: n=20 Veh. IBMX, n=20 IBMX, n=31 Veh. lidocain, n=37 lidocain, n=15 Veh. nifedipin, n=15 nifedipin, n=16 Veh. norepinephrine, n=21 norepinephrine. Arrest regarding the position (Fig 2D), n=59.
- 9hpi: n=22 Veh. lidocain, n=23 lidocain.
- 24hpi: n=24 Veh. IBXM, n=23 IBMX, n=20 Veh. lidocain, n=20 lidocain.
- 1675 at 3hpi, n=32; A431at 3hpi, n=37; JIMT1 at 3hpi, n=40; 4T1 at 3hpi, n=40; ZMEL1 at 3hpi, n=28.
- Lidocain starting 1hpi, 24hpi imaging, n=19.
- Veh. lidocain starting 1hpi, 24hpi imaging, n=16.

Exception: Fig1C-D (ZF whole embryo 3 hours post-injection). Sample size = 11 fish/3 independent experiments. Due to the size of the whole embryo, the imaging time and cost of acquisition are very high. Adding the fact that the results were straight-forward, we decided to limit the sample size.

Exception: Fig 2E (ZF injection with transfected siRNA cells). Because of the increased variability of the results, and the fact that all injected fish were take into account (including embryos bearing no arrested CTC), we reproduce the experiments five time. Total sample sizes are: n=77 siCtrl, n=63 siITGb1 #1, n=55 siITGb1 #2.

Zebrafish live injection (**Fig 3D,E,F,J,K,L**) was done in triplicate of independent injection batch of 5 fish. This experimental procedure asks fine settings and 5 min acquisition time following the injection. The following sample size are the “final number of fish / the number of events” (= cell or small cluster of 2 to 5 cells): n=7/54 Veh. IBMX, n=3/62 IBMX, n=4/42 vehicle Lidocain, n=8/46 lidocain.

Zebrafish live injection with 10μm beads, **Fig S2A**. 2 embryos showing more than 260 events in total were took into account for the quantification.

Zebrafish flow around arrested tumor cell study using 100 μm beads (**Fig 5L, Fig S6C**) is based on high-speed imaging: n=7 “open”, n=6 “partially closed” and n=6 “closed”.

Permeability assays in the treated zebrafish (**Fig S4D**) were done over 8 embryos per conditions.

Vessel diameter measurements (**Fig 2B, Fig S4A-C**) were done on 1 representative control embryo (Fig 2B) and 3 embryos per condition for the comparison of the effect of the treatments (Fig S4A-C).

For zebrafish electronic microscopy experiments (**Fig 2G, Fig 4K, Fig 5E**) sample size is 1. These experiments have the objective to be as descriptive as possible, reaching the best resolution existing, but not to be quantitative. The embryos, the region of interest and the timing was chosen during a dedicated injection experiment.

### Mouse experiments

For Mouse blood flow measurement (**Fig 7B**), results were obtained from more than 5,000 tumor cells from four different tumor cell lines (JIMT1br3, A2058br, PC14PE6br, H1). A total of 51-96 vessels from 12-13 regions (dimensions: 607.28x607.28x372.00 μm) of 3-6 animals per group were quantified. Box plots representing median values with 10th, 25th, 75th and 90th percentiles.

For experiments described in Fig.S7 (A-D), results were obtained from (A): Distance to first perfused vessels, n=19; (C): Dextran High, n=17 vessels. Dextran low, n=7 vessels; (D) Diameter, n= 98 vessels from 4 stacks.

### Microfluidic and other *in vitro* experiments

Microfluidic perfusion assays (**Fig 1G, Fig 2F, Fig S2B, Fig S4I, Fig 6D, E**) were reproduced 3 times, with independent HUVEC culture and independent siRNA transfection when used. For Fig 2F, n=62, 64 and 65 cells in siCtrl, siITGb1 #1 and siITGb1 #2 respectively.

For Fig 6E, quantification was done on n=67 cells in no flow and n=88 in flow conditions.

<u>Exception</u>: The data shown on the Fig 1G come from 6 independent experiments using 6-channels IBIDI μ-slides. During each experiments, all the perfusion conditions couldn’t not be realized each time, so we decided to increase the number of experiments, inducing a final sample size of 5 to 6 perfused channel per condition.

Live imaging of pCAG-EGFP transfected HUVEC (**Fig 6A**) was done on 4 independent experiments. In total, n=14 cells for the no flow condition and n=15 cells for the flow condition.

SEM experiment (**Fig 6B, D**) was repeated in two independent experiments. For the quantification in 6B, n=15 fields for the flow and n=10 fields for the no flow condition were quantified.

Flow study experiments of 100 μm beads around arrested CTCs in the microfluidic channels (**Fig 6C**) were performed during 2 independent experiments. Velocity analysis was made on 6 cells.

Microfluidic assays couple with optical trapping (**Fig S2C,D**) were repeated in three independent experiments, with independent HUVEC culture and independent siRNA transfection. Represented total sample sizes (number of trapped CTC) are n = 8 (37°) and n=6 (28°) (Fig S2C) and n=12, 13, 15 and n= 15, 23, 14 cells, respectively for adhesion and detachment experiments with siRNA Integrin cells (siCtrl, siITGb1#1 and siITGb1#2 – Fig S2D).

*In vitro* assays (adhesion and migration - **Fig S4E-G, Fig S5D and H**) was repeated three times independently. Furthermore, adhesion assay was done in triplicate.

Western blot analysis and immunofluorescence quantification (**Fig 2C, Fig S2E**) were obtained from three independent experiments as a validation for the siRNA efficiency.

Cell diameter measurement of CTCs from patients (**Fig 2A**) were done post-isolation from blood sample from 6 patients, resulting in n=2, 12, 1, 53, 10, 7 cells respectively. For 1675, D2A1, JIMT-1, A431, 4T1, ZMEL cells diameter, quantification was done using acquisitions of cells in suspension, n>90.

### Patient’s MRI for brain metastatic study (Fig 8A-D)

Number of BM for the 100 patients are provided for the entire cohort: BM originate from various tumor types including lung cancer (n=51), skin cancer (n=17), breast cancer (n=10), genitourinary (GU) cancer (n=10), gastrointestinal (GI) cancer (n=4), sarcoma (n=2) and cancer of unknown primary (n=6). The differences in CBF and MTT between all BM and BM-voxels were compared by the independent t-test. Voxels with more than one BM were weighted according to the number of BM occurring within the voxel. The paired samples t-test was used to compare the perfusion values and the volume of the BM.

## Data and software availability

For most of the analysis: GraphPad V5.04 to V6, PRISM – commercial dedicated software for statistical analysis.

For human patients data, statistical analysis was conducted using IBM SPSS Statistics^®^ software (IBM^®^ 2011, version 20, Armonk, New York, USA) and R (The R Foundation, version 3.3.1. Vienna, Austria)

Optical tweezers acquisitions were driven with Labview software and Nationnal Instruments acquisition boards. The signal analyses for the optical tweezers as for PIV were performed with IgorPro and ImageJ. The Michaelis Menten model was developed under Wolfram Mathematica. The heatmap reconstruction is a homewritten script running under Matlab (available to the scientific community).

The in silico vasculare architecture and numerical simulation were performed with with Freecad, Gmsh, AngioTK and Feel++ (See Resource table for more information). Probing of flow profiles around arrested tumor cells was performed using Imaris (Bitplane) as well as FIJI.

## Supplementary figure legends

**Figure S1. Related to Figure 1: CTCs arrest and extravasate in vascular regions with permissive flow profiles** (A) Experimental workflow. (B) Maximum projection of confocal stack of the zebrafish brain vasculature (green) containing extravasated TCs (red). (C) Heatmaps showing the adhesion pattern of 6 cell lines in the caudal plexus of the zebrafish embryos. (D) Confocal Z-projection of zebrafish posterior caudal plexus (CP) at 2 dpf. (E) Partial 3D segmentation (Amira) based on confocal acquisition (D). (F) Simplified AOC model of the zebrafish CP. (G) Image extracted from Movie S3, showing flow simulation results in the AOC model. (H) Velocity profiles obtained from flow simulation at several positions indicated in F. (I) Graphical representation of minimum, maximum and mean values of the blood flow velocity over the 5 positions from F.

**Figure S2. Related to Figure 2: Permissive flow profiles allow adhesion of CTCs to ECs** (A) Experimental set up, representative image and quantifications of the 10-μm beads injection. (B) Experimental set up and results of the adhesion efficacy regarding the temperature. (C) Experimental set up and quantification of the measured adhesion rupture force (n=8 and 6) of attached TC (top). Annotated image sequence from Movie 6 showing the detachment of a TC from the endothelial monolayer (bottom). (D) Experimental set up and quantification of the ratio of effective adhesion events upon OT (left) and OT-mediated detachment events (right) using siRNA-treated TCs. (E) Quantification of the Western blot analysis in Fig.2E and representative immunostaining images from siITGB1 cells.

**Figure S3. Related to Figure 3: Flow tuning *in vivo*** (A) Experimental workflow, and a representative image of the PIV analysis in the CP. Quantification of the duration of the flow under 400 μm/sec for each pharmacological condition in the dorsal aorta (position ISV 1, 4 and 8 from A) (n = 4 to 5 embryos per condition). (B) Experimental workflow of OT-mediated quantification of hemodynamic forces *in vivo* and annotated image showing trapped RBC in the DA (yellow) (see also Movie 11). Quantification and mapping of the flow force measured with OT on trapped RBCs for 3 conditions: lidocain, IBMX and control (breeding water). Values in A are mean ±SD. (C and E) Image extracted from Movie 10, showing flow simulation results in the AOC model for vehicles and lidocain (C) or IBMX (E). (D and F) Velocity profiles from flow simulation at several positions for each condition (as shown in Fig.S1F).

**Figure S4. Related to Figure 4: Blood flow tuning does not affect the architecture of the caudal plexus or tumor cells ability to migrate and adhere.** (A) 3D segmentation (IMOD) from one zebrafish posterior caudal plexus per condition (See also Movie 12). (B) Quantification of the total vascular surface from dorsal aorta (DA, red on A) and caudal veins (CV, blue on A) for each condition (n=3). (C) Quantification of the number of vascular loops for each condition (n=3). (D) Representative images of embryos injected with dextran 500kDa and imaged 3 hpi are shown (E) Representative images of wound closure for each condition 0, 9 and 24 hours after wound. (F) Quantification of the wound closure over time (n=3). (G) Representative image of TC stained with crystal violet during the adhesion test (left). Quantification of the absorbance measured at 590nm in the 4 conditions (right). (H) Quantification of the number of adhesion events per field after 10 min of perfusion of tumor cells in each condition using the microfluidic channels. (I) Quantification of the number of extravasated cells 16 hours post-perfusion of tumor cells (10min), with or without flow, in the microfluidic channels in each condition. *In vitro* experiment were done at least 3 times, in triplicate. Values in (B-C), (F-G) and (H,I) are mean ±SD.

**Figure S5. Related to Figure 4: Tuning PMA with two other drugs also perturb the behavior of CTCs in vivo**. (A) Quantification of the PMA with nifedipin (5μM) treatment compared to vehicle. (B) Number and location of arrested CTCs 3hpi using heatmapping, and quantification (C). (D) Adhesion and migration assay results of TCs assessed in presence of nifedipin. (E) Quantification of the PMA with norepinephrine (500μM) treatment compared to control. (F) Number and location of arrested CTCs 3hpi using heatmapping, and quantification (G). Data normalized to vehicle AVJ mean. (H) Adhesion and migration of TCs is assessed in presence of norepinephrine. (I) Heatmaps and quantification of the extravasation efficacy of 1675, JIMT-1 and ZMEL cells 24 hpi. Values in (A), (C-F), (H-J) and (I) are mean ±SD.

**Figure S6. Related to Figure 5L: Flow profiles around (and close to) arrested tumor cells in the zebrafish embryo** (A) Scheme depicting flow profiles around arrested TC in the zebrafish embryo (Open, partially clogged and clogged) and quantification of the distance between arrested TC and the first fully perfused vessel (B) Images of *in silico* simulations for the respective scenario (C) Images and analysis of high-speed confocal acquisitions in the zebrafish embryo. Values in (A) are mean ±SD.

**Figure S7. Related to Figure 7: Arrested CTCs in the mouse brain are in contact (or close to) with laminar flow profiles**. (A) Scheme depicting flow profiles around arrested TC in the mouse brain and quantification of the distance between arrested TC and the first fully perfused vessel. (B) 3D image of an arrested TC in a cerebral capillary. (C) Quantification of Dextran intensity before and after the arrested TC, ratio is plotted for both High (HMW) and Low (LMW) Molecular Weight Dextran. (D) Quantification of mean vessel diameter in vessel bearing (with TC) or not (neighboring) arrested TC. Values in (A), (C), and (D) are mean ±SD.

## Supplementary Movie Legends

**Movie S1. Related to Figure 1: CTCs arrest and extravasate in the brain of zebrafish embryos**. Low (top-left) and high-magnification (right) confocal Z-stack of the zebrafish head vasculature (green), containing two extravasated TCs (red). (bottom-left) high-speed imaging of the blood flow in the vessel of extravasated TCs.

**Movie S2. Related to Figure 1: Blood flow drops along the arterial vasculature of the zebrafish embryo**. Whole-embryo high-speed flow acquisition and PIV analysis. A selection of 4 regions of interest reveal the significant flow drop in the caudal region. Color code = velocity ranging from 2500 (red) to 50 μm/sec.

**Movie S3. Related to Figure S1, related to Figure 1: Flow drop in the caudal plexus occurs *in silico***. 3D reconstruction of the caudal plexus is used for blood flow simulation.

**Movie S4. Related to Figure 2: Fine vascular architecture of a 2-dpf zebrafish embryo**. 3D reconstruction of the vascular architecture of a 2-dpf zebrafish embryo, upon two-photon excitation microscopy.

**Movie S5. Related to Figure 2: Stable adhesion and intravascular migration of CTCs**. Time-lapse analysis of the intravascular adhesion of TCs (red) to the endothelium (green) of 2-dpf embryos *(Tg(fli1a:egfp)),* in the caudal plexus. Two different z position are provided.

**Movie S6. Related to Figure S2, related to Figure 2: Identification of early adhesion forces of CTCs to ECs**. Transmitted light imaging of OT experiments *in vitro* shows the adhesion and its rupture between a single CTC (trapped) and ECs. The plot displays the extracted adhesion resistance force for this event. Adhesion to EC imposes a growing force (25-40 sec) before its break, visible on the graph with the drop of force measured (40-45 sec).

**Movie S7. Related to Figure S2, related to Figure 2: Depletion of ITGB1 impedes the early adhesion of CTCs to ECs**. Transmitted light imaging of optical trapping of siCTL versus siITGB1 cells. Stable adhesion of the CTC to the EC monolayer is visible in the control situation (See time=20s). No adhesion is visible for siITGB1 cells.

**Movie S8. Related to Figure 2: Early adhesion forces of CTCs to ECs *in vivo* rapidly exceed 200 pN**. X,Y and Z movement of the optical trap *in vivo* has no effect on arrested CTCs in the dorsal aorta. Successful trapping of RBCs in close contact with the arrested CTC, which remains attached to ECs, shows that adhesion forces rapidly exceed 200 pN *in vivo.*

**Movie S9. Related to Figure 3: Tuning the blood flow in the CP of 2-dpf zebrafish embryos**. High-speed flow acquisition (200 fps) and PIV analysis in the CP of vehicle- or drug-treated zebrafish embryos. Color code = velocity ranging from 1600 (red) to 400 μm/sec.

**Movie S10. Related to Figure S3, related to Figure 3: In silico tuning of the blood flow in the virtual CP**. 3D reconstruction of the CP is used for blood flow simulation. Flow frequencies measured *in vivo* are used (values in brackets).

**Movie S11. Related to Figure S3, related to Figure 3: Tuning and measuring hemodynamic forces *in vivo***. Transmitted light imaging of OT of RBCs performed in the DA of zebrafish embryos. Movie(s) show efficient trapping of RBCs in the DA and the CV. Red circle = OT.

**Movie S12. Related to Figure S4, related to Figure 3: Tuning the blood flow does not perturb the vascular architecture of the CP**. 3D segmentations of the caudal plexus in the 4 different conditions are shown. Dorsal aorta appears in red, caudal veins in blue.

**Movie S13. Related to Figure 3: Tuning blood flow forces perturbs the early arrest of CTCs in living zebrafish embryos**. Instantaneous live imaging of CTCs (red) upon intravascular (green) injection in vehicle- or drug-treated zebrafish embryos reveals that blood flow affects the early arrest of CTCs. Final Color code is based on time of arrival and residency of tumor cells (Image J).

**Movie S14. Related to Figure 5: CLEM of extravasated TCs in the zebrafish embryo**. Confocal imaging and correlative serial TEM of TCs reveal the extra- and intra-vascular location.

**Movie S15. Related to Figure 5: Extravasation of arrested CTCs in the caudal plexus of a zebrafish embryo**. 3D time-lapse confocal imaging of arrested CTCs (red) in the CP (green) over 15h. Extravasation of CTCs occurs mostly upon ECs pocketing and remodeling.

**Movie S16. Related to Figure 5 Vascular pocketing and remodeling drives extravasation of arrested CTCs**. Confocal imaging, correlative serial TEM and electron tomography of extravasating TCs (red) shows discrete pocketing by ECs (green).

**Movie S17. Related to Figure 6: Flow forces stimulate protrusive activity of the dorsal surface of endothelial cells**. Confocal time-lapse imaging of two endothelial cells, transfected with pCAG-EGFP, stimulated with laminar flows (400 μm/s).

## REFERENCES

1. Nguyen, D. X., Bos, P. D. & Massagué, J. Metastasis: from dissemination to organ-specific colonization. Nat. Rev. Cancer 9, 274–284 (2009).

2. Hosseini, H. et al. Early dissemination seeds metastasis in breast cancer. Nature (2016). doi:10.1038/nature20785

3. Harper, K. L. et al. Mechanism of early dissemination and metastasis in Her2(+) mammary cancer. Nature (2016). doi:10.1038/nature20609

4. Valastyan, S. & Weinberg, R. A. Tumor Metastasis: Molecular Insights and Evolving Paradigms. Cell 147, 275–292 (2011).

5. Regmi, S., Fu, A. & Luo, K. Q. High Shear Stresses under Exercise Condition Destroy Circulating Tumor Cells in a Microfluidic System. Scientific Reports 7, 39975 (2017).

6. Sosa, M. S., Bragado, P. & Aguirre-Ghiso, J. A. Mechanisms of disseminated cancer cell dormancy: an awakening field. Nat. Rev. Cancer 14, 611–622 (2014).

7. Kienast, Y. et al. Real-time imaging reveals the single steps of brain metastasis formation. Nat. Med. 16, 116–122 (2010).

8. Chen, Q. et al. Carcinoma-astrocyte gap junctions promote brain metastasis by cGAMP transfer. Nature 533, 493–498 (2016).

9. Headley, M. B. et al. Visualization of immediate immune responses to pioneer metastatic cells in the lung. Nature 531, 513–517 (2016).

10. Strilic, B. et al. Tumour-cell-induced endothelial cell necroptosis via death receptor 6 promotes metastasis. Nature 536, 215–218 (2016).

11. Paszek, M. J. et al. Tensional homeostasis and the malignant phenotype. Cancer Cell 8, 241–254 (2005).

12. Levental, K. R. et al. Matrix crosslinking forces tumor progression by enhancing integrin signaling. Cell 139, 891–906 (2009).

13. Mouw, J. K. et al. Tissue mechanics modulate microRNA-dependent PTEN expression to regulate malignant progression. Nat. Med. 20, 360–367 (2014).

14. Stylianopoulos, T. et al. Causes, consequences, and remedies for growth-induced solid stress in murine and human tumors. Proc. Natl. Acad. Sci. U.S.A. 109, 15101–15108 (2012).

15. Chauhan, V. P. et al. Angiotensin inhibition enhances drug delivery and potentiates chemotherapy by decompressing tumour blood vessels. Nat Commun 4, 2516 (2013).

16. Provenzano, P. P. et al. Enzymatic targeting of the stroma ablates physical barriers to treatment of pancreatic ductal adenocarcinoma. Cancer Cell 21, 418–429 (2012).

17. Wolf, K. et al. Physical limits of cell migration: control by ECM space and nuclear deformation and tuning by proteolysis and traction force. J. Cell Biol. 201, 1069–1084 (2013).

18. Harada, T. et al. Nuclear lamin stiffness is a barrier to 3D migration, but softness can limit survival. J. Cell Biol. 204, 669–682 (2014).

19. Denais, C. M. et al. Nuclear envelope rupture and repair during cancer cell migration. Science 352, 353–358 (2016).

20. Irianto, J. et al. DNA Damage Follows Repair Factor Depletion and Portends Genome Variation in Cancer Cells after Pore Migration. Curr. Biol. (2016). doi:10.1016/j.cub.2016.11.049

21. Paget, S. The distribution of secondary growths in cancer of the breast. 1889. Cancer Metastasis Rev. 8, 98–101 (1989).

22. Weiss, L. Comments on hematogenous metastatic patterns in humans as revealed by autopsy. Clin. Exp. Metastasis 10, 191–199 (1992).

23. Chambers, A. F., Groom, A. C. & MacDonald, I. C. Dissemination and growth of cancer cells in metastatic sites. Nat. Rev. Cancer 2, 563–572 (2002).

24. Labelle, M., Begum, S. & Hynes, R. O. Platelets guide the formation of early metastatic niches. Proc. Natl. Acad. Sci. U.S.A. 111, E3053–3061 (2014).

25. Chang, S.-F. et al. Tumor cell cycle arrest induced by shear stress: Roles of integrins and Smad. PNAS 105, 3927–3932 (2008).

26. Wirtz, D., Konstantopoulos, K. & Searson, P. C. The physics of cancer: the role of physical interactions and mechanical forces in metastasis. Nat. Rev. Cancer 11, 512–522 (2011).

27. Follain, G., Mercier, L., Osmani, N., Harlepp, S. & Goetz, J. G. Seeing is believing-multi-scale spatio-temporal imaging towards in vivo cell biology. J Cell Sci 130, 23–38 (2017).

28. Karreman, M. A., Hyenne, V., Schwab, Y. & Goetz, J. G. Intravital Correlative Microscopy: Imaging Life at the Nanoscale. Trends Cell Biol. 26, 848–863 (2016).

29. Karreman, M. A. et al. Fast and precise targeting of single tumor cells in vivo by multimodal correlative microscopy. J Cell Sci 129, 444–456 (2016).

30. Stoletov, K. et al. Visualizing extravasation dynamics of metastatic tumor cells. J Cell Sci 123, 2332–2341 (2010).

31. Anton, H. et al. Pulse propagation by a capacitive mechanism drives embryonic blood flow. Development 140, 4426–4434 (2013).

32. Reymond, N., d’Água, B. B. & Ridley, A. J. Crossing the endothelial barrier during metastasis. Nat. Rev. Cancer 13, 858–870 (2013).

33. Cameron, M. D. et al. Temporal progression of metastasis in lung: cell survival, dormancy, and location dependence of metastatic inefficiency. Cancer research 60, 2541–2546 (2000).

34. Luzzi, K. J. et al. Multistep nature of metastatic inefficiency: dormancy of solitary cells after successful extravasation and limited survival of early micrometastases. The American journal of pathology 153, 865–873 (1998).

35. Seguin, L., Desgrosellier, J. S., Weis, S. M. & Cheresh, D. A. Integrins and cancer: regulators of cancer stemness, metastasis, and drug resistance. Trends Cell Biol. 25, 234–240 (2015).

36. Gassmann, P., Hemping-Bovenkerk, A., Mees, S. T. & Haier, J. Metastatic tumor cell arrest in the liver-lumen occlusion and specific adhesion are not exclusive. International Journal of Colorectal Disease 24, 851–858 (2009).

37. Schlesinger, M. & Bendas, G. Contribution of very late antigen-4 (VLA-4) integrin to cancer progression and metastasis. Cancer Metastasis Rev. 34, 575–591 (2015).

38. Goetz, J. G. et al. Endothelial Cilia Mediate Low Flow Sensing during Zebrafish Vascular Development. Cell Reports 6, 799–808 (2014).

39. Vermot, J. et al. Reversing blood flows act through klf2a to ensure normal valvulogenesis in the developing heart. PLoS Biol. 7, e1000246 (2009).

40. Luca, E. D. et al. ZebraBeat: a flexible platform for the analysis of the cardiac rate in zebrafish embryos. Scientific Reports 4, 4898 (2014).

41. Brennan, C. Acetylcholine and calcium signalling regulates muscle fibre formation in the zebrafish embryo. Journal of Cell Science 118, 5181–5190 (2005).

42. Kai, F., Laklai, H. & Weaver, V. M. Force Matters: Biomechanical Regulation of Cell Invasion and Migration in Disease. Trends Cell Biol. 26, 486–497 (2016).

43. Azevedo, A. S., Follain, G., Patthabhiraman, S., Harlepp, S. & Goetz, J. G. Metastasis of circulating tumor cells: favorable soil or suitable biomechanics, or both? Cell Adh Migr 9, 345–356 (2015).

44. Heidemann, F. et al. Selectins mediate small cell lung cancer systemic metastasis. PLoS ONE 9, e92327 (2014).

45. Cheung, K. J. et al. Polyclonal breast cancer metastases arise from collective dissemination of keratin 14-expressing tumor cell clusters. Proc. Natl. Acad. Sci. U.S.A. 113, E854–863 (2016).

46. Cheung, K. J. & Ewald, A. J. A collective route to metastasis: Seeding by tumor cell clusters. Science 352, 167–169 (2016).

47. Aceto, N. et al. Circulating tumor cell clusters are oligoclonal precursors of breast cancer metastasis. Cell 158, 1110–1122 (2014).

48. Hwang, T. L., Close, T. P., Grego, J. M., Brannon, W. L. & Gonzales, F. Predilection of brain metastasis in gray and white matter junction and vascular border zones. Cancer 77, 1551–1555 (1996).

49. Chen, M. B., Lamar, J. M., Li, R., Hynes, R. O. & Kamm, R. D. Elucidation of the Roles of Tumor Integrin β1 in the Extravasation Stage of the Metastasis Cascade. Cancer Res. 76, 2513–2524 (2016).

50. Klemke, M., Weschenfelder, T., Konstandin, M. H. & Samstag, Y. High affinity interaction of integrin alpha4beta1 (VLA-4) and vascular cell adhesion molecule 1 (VCAM-1) enhances migration of human melanoma cells across activated endothelial cell layers. J. Cell. Physiol. 212, 368–374 (2007).

51. Glinskii, O. V. et al. Mechanical entrapment is insufficient and intercellular adhesion is essential for metastatic cell arrest in distant organs. Neoplasia 7, 522–527 (2005).

52. Enns, A. et al. Integrins can directly mediate metastatic tumor cell adhesion within the liver sinusoids. J. Gastrointest. Surg. 8, 1049–1059; discussion 1060 (2004).

53. Petri, B. et al. Endothelial LSP1 is involved in endothelial dome formation, minimizing vascular permeability changes during neutrophil transmigration in vivo. Blood 117, 942–952 (2011).

54. Phillipson, M., Kaur, J., Colarusso, P., Ballantyne, C. M. & Kubes, P. Endothelial domes encapsulate adherent neutrophils and minimize increases in vascular permeability in paracellular and transcellular emigration. PLoS ONE 3, e1649 (2008).

55. Carman, C. V. & Springer, T. A. A transmigratory cup in leukocyte diapedesis both through individual vascular endothelial cells and between them. J. Cell Biol. 167, 377–388 (2004).

56. Allen, T. A. et al. Angiopellosis as an Alternative Mechanism of Cell Extravasation. Stem Cells 35, 170–180 (2017).

57. Heyder, C. et al. Realtime visualization of tumor cell/endothelial cell interactions during transmigration across the endothelial barrier. J. Cancer Res. Clin. Oncol. 128, 533–538 (2002).

58. Khuon, S. et al. Myosin light chain kinase mediates transcellular intravasation of breast cancer cells through the underlying endothelial cells: a three-dimensional FRET study. J Cell Sci 123, 431–440 (2010).

59. Tremblay, P.-L., Huot, J. & Auger, F. A. Mechanisms by which E-selectin regulates diapedesis of colon cancer cells under flow conditions. Cancer Res. 68, 5167–5176 (2008).

60. Al-Mehdi, A. B. et al. Intravascular origin of metastasis from the proliferation of endothelium-attached tumor cells: a new model for metastasis. Nat Med 6, 100–102 (2000).

61. Lapis, K., Paku, S. & Liotta, L. A. Endothelialization of embolized tumor cells during metastasis formation. Clinical & experimental metastasis 6, 73–89 (1988).

62. Lambert, A. W., Pattabiraman, D. R. & Weinberg, R. A. Emerging Biological Principles of Metastasis. Cell 168, 670–691 (2017).

63. Grutzendler, J. et al. Angiophagy prevents early embolus washout but recanalizes microvessels through embolus extravasation. Sci Transl Med 6, 226ra31 (2014).

64. Lam, C. K., Yoo, T., Hiner, B., Liu, Z. & Grutzendler, J. Embolus extravasation is an alternative mechanism for cerebral microvascular recanalization. Nature 465, 478–482 (2010).

65. Franco, C. A. et al. Dynamic endothelial cell rearrangements drive developmental vessel regression. PLoS Biol. 13, e1002125 (2015).

66. Gebala, V., Collins, R., Geudens, I., Phng, L.-K. & Gerhardt, H. Blood flow drives lumen formation by inverse membrane blebbing during angiogenesis in vivo. Nat. Cell Biol. 18, 443–450 (2016).

67. Potente, M., Gerhardt, H. & Carmeliet, P. Basic and therapeutic aspects of angiogenesis. Cell 146, 873–887 (2011).

68. Osswald, M. et al. Brain tumour cells interconnect to a functional and resistant network. Nature 528, 93–98 (2015).

69. Shibue, T., Brooks, M. W. & Weinberg, R. A. An integrin-linked machinery of cytoskeletal regulation that enables experimental tumor initiation and metastatic colonization. Cancer Cell 24, 481–498 (2013).

70. Hvichia, G. E. et al. A novel microfluidic platform for size and deformability based separation and the subsequent molecular characterization of viable circulating tumor cells. Int. J. Cancer 138, 2894–2904 (2016).

71. Chudziak, J. et al. Clinical evaluation of a novel microfluidic device for epitope-independent enrichment of circulating tumour cells in patients with small cell lung cancer. Analyst 141, 669–678 (2016).

72. Magbanua, M. J. M. et al. A Novel Strategy for Detection and Enumeration of Circulating Rare Cell Populations in Metastatic Cancer Patients Using Automated Microfluidic Filtration and Multiplex Immunoassay. PLoS ONE 10, e0141166 (2015).

73. Robb, R. A. The biomedical imaging resource at Mayo Clinic. IEEE Trans Med Imaging 20, 854–867 (2001).

74. Nudelman, K. N. H. et al. Altered cerebral blood flow one month after systemic chemotherapy for breast cancer: a prospective study using pulsed arterial spin labeling MRI perfusion. PLoS ONE 9, e96713 (2014).

75. Schwarzbach, C. J. et al. Stroke and cancer: the importance of cancer-associated hypercoagulation as a possible stroke etiology. Stroke 43, 3029–3034 (2012).

76. Feyen, L. et al. Standardization of Dynamic Whole-Brain Perfusion CT: A Comprehensive Database of Regional Perfusion Parameters. 189–220 (2010).

77. Abels, B., Klotz, E., Tomandl, B. F., Kloska, S. P. & Lell, M. M. Perfusion CT in Acute Ischemic Stroke: A Qualitative and Quantitative Comparison of Deconvolution and Maximum Slope Approach. American Journal of Neuroradiology 31, 1690–1698 (2010).

78. Kemmling, A. et al. Decomposing the Hounsfield unit: probabilistic segmentation of brain tissue in computed tomography. Clin Neuroradiol 22, 79–91 (2012).

